# Reflectance spectroscopy allows rapid, accurate, and non-destructive estimates of functional traits from pressed leaves

**DOI:** 10.1101/2021.04.21.440856

**Authors:** Shan Kothari, Rosalie Beauchamp-Rioux, Etienne Laliberté, Jeannine Cavender-Bares

## Abstract

1. More than ever, ecologists seek to employ herbarium collections to estimate plant functional traits from the past and across biomes. However, many trait measurements are destructive, which may preclude their use on valuable specimens. Researchers increasingly use reflectance spectroscopy to estimate traits from fresh or ground leaves, and to delimit or identify taxa. Here, we extend this body of work to non-destructive measurements on pressed, intact leaves, like those in herbarium collections.
2. Using 618 samples from 68 species, we used partial least-squares regression to build models linking pressed-leaf reflectance spectra to a broad suite of traits, including leaf mass per area (LMA), leaf dry matter content (LDMC), equivalent water thickness, carbon fractions, pigments, and twelve elements. We compared these models to those trained on fresh- or ground-leaf spectra of the same samples.
3. Our pressed-leaf models were best at estimating LMA (*R^2^* = 0.932; %RMSE = 6.56), C (*R^2^* = 0.855; %RMSE = 9.03), and cellulose (*R^2^* = 0.803; %RMSE = 12.2), followed by water-related traits, certain nutrients (Ca, Mg, N, and P), other carbon fractions, and pigments (all *R^2^* = 0.514-0.790; %RMSE = 12.8-19.6). Remaining elements were predicted poorly (*R^2^* < 0.5, %RMSE > 20). For most chemical traits, pressed-leaf models performed better than fresh-leaf models, but worse than ground-leaf models. Pressed-leaf models were worse than fresh-leaf models for estimating LMA and LDMC, but better than ground-leaf models for LMA. Finally, in a subset of samples, we used partial least-squares discriminant analysis to classify specimens among 10 species with near-perfect accuracy (>97%) from pressed- and ground-leaf spectra, and slightly lower accuracy (>93%) from fresh-leaf spectra.
4. These results show that applying spectroscopy to pressed leaves is a promising way to estimate leaf functional traits and identify species without destructive analysis. Pressed-leaf spectra might combine advantages of fresh and ground leaves: like fresh leaves, they retain some of the spectral expression of leaf structure; but like ground leaves, they circumvent the masking effect of water absorption. Our study has far-reaching implications for capturing the wide range of functional and taxonomic information in the world’s preserved plant collections.

## Introduction

The world’s herbaria together contain more than 390 million specimens (Thiers 2021) which are a rich source of information about global plant diversity. Herbarium specimens are collected for many reasons—often to document where a species is present, or to serve as vouchers for taxonomic studies. But these specimens are often repurposed for new ends, unforeseen by their collectors (Meineke et al. 2018). More than ever, ecologists and evolutionary biologists seek to use herbarium specimens to measure functional traits (Heberling 2021): for example, to evaluate the long-term imprint of human activity on plant communities (Lang et al. 2018; Meineke et al. 2018); to fill in gaps in sparse trait databases (Perez et al. 2020); or to conduct comparative studies of clades (Jardine et al. 2020). Measuring functional traits on herbarium specimens carries the promise of letting us reach the inaccessible, including the past or distant parts of the world. Using herbarium specimens also allows researchers to benefit from the expertise of taxonomists and refer back to the same specimens for further use—for example, as sources of genetic data, or as references for species identification from new collections (Heberling 2021). Using specimens, researchers can address ecological and evolutionary questions that require merging functional, genetic, and distributional data at global scales.

Many functional trait measurements require destructive sampling—for example, by grinding up tissue for chemical analyses. Such measurements include most protocols to determine the elemental or molecular composition of a sample (Pérez-Harguindeguy et al. 2013). Because herbarium specimens are irreplaceable—especially those from historical collections—curators may hesitate to let them be destroyed, even in part, for ecological research. Using specimens in functional ecology might be more feasible with new, non-destructive techniques to estimate their traits.

Reflectance spectroscopy is a technique often used to estimate foliar functional traits non-destructively (Curran 1989; Jacquemoud & Ustin 2020). Spectroscopy is the study of matter’s interactions with electromagnetic radiation across wavelengths (Jacquemoud & Ustin 2020); spectroscopic studies of leaves often target reflectance—the proportion of incident radiation that is reflected—as a particularly revealing and easy-to-measure property. A typical leaf reflectance spectrum comprises reflectance at many narrow wavelength bands between about 350 and 2500 nm, which includes over 98% of energy from solar radiation reaching Earth’s surface (American Society for Testing and Materials, 2006). Because the leaf’s chemical and structural makeup determines how it reflects, absorbs, and transmits light, reflectance within this range carries information about many plant traits (Cavender-Bares et al. 2017).

Two main approaches exist to estimate traits using the full information in reflectance spectra. First, physics-based radiative transfer models like PROSPECT can be inverted to estimate a handful of traits with well-defined optical properties (Féret et al. 2017). Second, statistical models, often created using machine learning techniques like partial least-squares regression (PLSR), can estimate an even wider range of traits, albeit in a less mechanistic (and perhaps less general) way (Serbin & Townsend 2020). This multivariate empirical approach gives researchers the flexibility to predict complex traits whose absorption features might not be as strong or well-defined (Curran 1989). Likewise, multivariate classification techniques like partial least-squares discriminant analysis (PLS-DA) use the full spectrum to discriminate species, lineages, or other kinds of biological classes (Meireles et al. 2020b).

Empirical approaches like PLSR are widely used to estimate plant traits from spectroscopic data measured on fresh or ground leaves. These traits include leaf N and leaf mass per area (LMA; Asner et al. 2011; Serbin et al. 2014; Serbin et al. 2019; Streher et al. 2020), pigments (Asner et al. 2011; Yang et al. 2016), defense compounds (Couture et al. 2016; Nakaji et al. 2019), non-structural carbohydrates (Ely et al. 2019), and even photosynthetic capacity (Yan et al. 2021). Leaf-level PLSR models have been used to address such varied ecological topics as defense responses to herbivory (Kula et al. 2020) and the role of biodiversity in ecosystem function (Schweiger et al. 2018). Although this multivariate statistical approach is flexible, it is sensitive to the kind of leaf tissue used to train the model. Existing PLSR models have mostly been trained on reflectance spectra of fresh leaves (e.g. Serbin et al. 2019) or dried, ground leaves (e.g. Serbin et al. 2014). Such models are not expected to transfer to dried, intact leaves like herbarium specimens because both drying and grinding cause major changes in reflectance.

We built PLSR models to estimate traits from the reflectance spectra of pressed leaves, like herbarium specimens, and compared their accuracy to models built from fresh or dried, ground leaves. Our pressed leaf samples were prepared like herbarium specimens but not yet mounted on paper, and our analyses might thus serve as a proof of concept for the technique by setting aside the methodological challenges related to working with older mounted specimens. Previously, Costa et al. (2018) had shown that a related spectroscopic technique, Fourier Transform-Near Infrared Spectroscopy (FT-NIR), could predict several leaf structural traits from pressed leaves of tropical trees. Here, we explicitly compare the accuracy of trait estimation from the spectra of fresh, pressed, and ground leaves for many leaf chemical and structural traits from multiple biomes and functional groups.

For most chemical traits, such as elemental composition or carbon fractions, we conjectured that ground-leaf spectral models would be the most accurate, as others have found (Serbin et al. 2014; Couture et al. 2016; Wang et al. 2020). Both drying and grinding might be important in achieving this accuracy. Drying may reveal minor absorption features of compounds in dry matter within the short-wave infrared (SWIR) range that, in fresh leaves, are obscured by the dominant effect of water absorption (Peterson et al. 1988). Grinding homogenizes the variation in structure and composition throughout the leaf lamina, which may allow us to capture a more even and representative sample of tissue (Richardson et al. 2021). Since pressed leaves are dried but not ground, we predicted they would yield intermediate accuracy for chemical traits.

For structural traits like LMA, we instead expected fresh and pressed leaves to outperform ground leaves because grinding disrupts the leaf structure. For water-related traits like leaf dry matter content (LDMC; dried mass divided by fresh mass), we expected fresh leaves to outperform both pressed and ground leaves because they retain the water absorption features that allow direct prediction of water content (Carter 1991). Likewise, because pigments tend to degrade after collection, we expected to estimate their concentration best from fresh leaves. We also assessed sample discoloration and considered whether it reduces the accuracy of trait estimates, which may indicate whether spectroscopy is useful on old or degraded specimens.

Finally, we asked whether pressed-leaf spectra can be used to identify samples to species. Reflectance spectra often show phylogenetic signal in certain wavelength ranges because of phylogenetic conservatism in their underlying traits (McManus et al. 2016; Meireles et al. 2020a; Diniz et al. 2020). This signal is what often makes it possible to classify species or higher-level taxa from fresh-leaf spectra (Cavender-Bares et al. 2016; Meireles et al. 2020b). Studies with tropical forest species have also shown that FT-NIR absorbance spectra of pressed leaves can be used to classify species or higher-level taxa (Durgante et al. 2013; Lang et al. 2015; Prata et al. 2018). Based on these studies, Draper et al. (2020) proposed using spectra of herbarium specimens as part of an integrative process of species delimitation and identification. However, it remains uncertain whether fresh- or pressed-leaf spectra are better suited to the task of classifying species. Here, we compared the accuracy of supervised classification from fresh-, pressed-, and ground-leaf spectra among the common species in our dataset. Because they preserve some of the leaf structure (unlike ground leaves) and reveal the distinctive SWIR absorption features of macromolecules and other compounds (unlike fresh leaves), pressed leaves may represent the best of both worlds for distinguishing species using spectroscopy.

## Methods

### Spectral and leaf trait measurements

We trained PLSR models on leaf reflectance spectra and traits measured as part of four projects conducted by the Canadian Airborne Biodiversity Observatory (CABO). We validated the models both internally and on an independent dataset of pressed tree and herb samples collected at Cedar Creek Ecosystem Science Reserve (East Bethel, MN, USA). Table 1 describes the projects and lists how many samples and species they include for each functional group. The leaf sampling procedure is described in the Supplementary Materials.

**Table 1:** A summary of CABO projects used for model building and internal validation (above the thick black line) and the Cedar Creek dataset used for external validation (below the thick black line). The column heading ‘broadleaf’ refers to broadleaf trees only.

|  |  | Samples (species) per functional group |  |  |  |  |  |
| --- | --- | --- | --- | --- | --- | --- | --- |
| Project | Description | Broadleaf | Conifer | Herb | Shrub | Vine | Total |
| Beauchamp-Rioux | Deciduous forest trees sampled throughout the growing season in 2018 at several sites in southern Québec and Ontario | 405 (9) |  |  |  |  | 405 (9) |
| Dessain | Forbs, grasses, shrubs, and broadleaf trees from forests and open areas sampled throughout the growing season in 2017 at four sites in southern Québec | 31 (24) |  | 15 (13) | 26 (22) |  | 72 (59) |
| Boucherville | Forbs, shrubs, and grasses sampled throughout the growing season in 2018 from the Parc national des Îles-de-Boucherville in southern Québec |  |  | 45 (8) | 22 (3) | 6 (1) | 73 (12) |
| Warren | A single broadleaf evergreen tree species (*Agonis flexuosa* (Willd.) Sweet) collected across a soil chronosequence (Turner et al. 2018) in southwestern Australia | 68 (1) |  |  |  |  | 68 (1) |
| Total |  | 504 (25) |  | 60 (19) | 48 (23) | 6 (1) | 618 (68) |
| Cedar Creek | Herbs and broadleaf and coniferous trees sampled at Cedar Creek LTER in Minnesota in late summer and fall 2018 | 177 (10) | 101 (4) | 55 (14) |  |  | 333 (28) |

For each CABO sample, we measured full-range reflectance spectra (350-2500 nm) of the leaves at three stages: (1) freshly sampled, (2) pressed and (3) oven-dried and ground into a fine powder. We measured fresh-leaf directional-hemispherical reflectance spectra on the adaxial surface of multiple leaves or leaf arrays from each sample using spectroradiometers equipped with integrating spheres. We pressed a portion of the sample, and measured reflectance spectra on the adaxial surface of pressed leaves between six months and three years later using a spectroradiometer with a leaf clip (Fig. S1). Lastly, we measured ground-leaf spectra using a spectroradiometer with a benchtop reflectance probe that pressed loose leaf powder into an even pellet with very low transmittance. We trimmed all spectra to 400-2400 nm. Detailed notes on measurement and post-measurement processing of reflectance spectra are found in the Supplementary Materials.

While measuring pressed-leaf spectra, we inspected each pressed specimen by eye to note signs of discoloration in preparation or storage. While all leaves have some changes in their appearance as they dry, we were particularly interested in the loss of green color, such as blackening, browning, or the development of a silvery or whitish finish on the leaf surface. We scored each leaf on a discrete scale from 0 to 4 (see examples in Figs. S2-5). A score of 0 indicates no noticeable discoloration. Scores 1 through 4 indicate increasing discoloration, from 1 (either <10% blackening/browning or development of a slight silvery finish to the leaf) to 4 (>75% blackening/browning).

We measured the following leaf structural and chemical traits on each CABO sample: Leaf mass per area (LMA; kg m^-2^), leaf dry matter content (LDMC; mg g^-1^), equivalent water thickness (EWT; mm), carbon fractions (soluble cell contents, hemicellulose, cellulose, and lignin; %), pigments (chlorophyll *a*, chlorophyll *b*, and total carotenoids; mg g^-1^), and concentrations of a variety of elements (Al, C, Ca, Cu, Fe, K, Mg, Mn, N, Na, P, Zn; % or mg g^-1^). Protocol summaries are in the Supplementary Materials.

### PLSR modeling for trait estimation

We used a PLSR modeling framework to predict each trait from each of fresh-, pressed-, and ground-leaf spectra across the full range (400-2400 nm). PLSR is suited to handle spectral datasets, which have many collinear predictors, because it projects the spectral matrix onto a smaller number of orthogonal latent components in a way that maximizes the ability to predict the response variable. We simply used reflectance values as predictors, since calculating various common transformations of reflectance (see Serbin et al. 2014) did not increase predictive accuracy in preliminary tests. We also did not transform any trait values to reduce skewness, again finding in preliminary tests that it did not improve predictive accuracy enough to warrant the added complexity. Some studies restrict the wavelengths used in prediction, often to ranges known or assumed to contain features relevant to a given trait (e.g. Serbin et al. 2014). We considered that the 400-1300 nm range might be most liable to change in storage due to degradation of photosynthetic pigments and accumulation of brown pigments that absorb in the near-infrared range (NIR; Fourty et al. 1996). Thus, we also built pressed-leaf models restricted to 1300-2400 nm (‘restricted-range models’), which might be expected to generalize better to datasets that include older or more discolored leaves. We present these results mainly in the Supplementary Materials.

Our methods for model calibration and validation largely follow Burnett et al. (2021). First, we randomly divided the data into calibration (75%) and validation (25%) datasets, stratified by functional group. We began by fitting a model for each trait on the calibration dataset. We selected the smallest number of components for which the root mean squared error of prediction (RMSEP) from 10-fold cross-validation fell within one standard deviation of the global minimum. We used this number of components—a different number for each trait—in further analyses to predict traits on the internal validation dataset. We calculated the variable importance in projection (VIP) metric for calibration models to see which parts of the spectrum were most important for predicting each trait (Wold et al. 1994).

To test how well we could predict traits on the internal validation subset, we first did a jackknife analysis by iteratively (100×) dividing the 75% calibration data further into random 70% training and 30% testing subsets. For each trait, we trained models on the 70% using the previously determined optimal number of components and predicted the remaining 30%. This analysis gave us a distribution of model performance statistics across the 100 iterates (*R^2^*, %RMSE), which reveals the sensitivity of model performance to randomly varying sets of training and testing data.

Next, we applied the 100 jackknife models for each trait to the 25% internal validation dataset, yielding a distribution of 100 predictions for each validation sample. We quantified model performance using *R^2^* and root mean squared error (RMSE) between measurements and mean predictions. We also report the RMSE as a percentage of the 2.5% trimmed range of measured values (%RMSE), which we used rather than the entire range (as in e.g. Burnett et al. 2021) for robustness to outliers. For each trait, we also tested whether the magnitude of residuals (observed minus predicted) in the validation dataset varied among leaves with different discoloration scores. We performed all statistical analyses in *R v. 3.6.3* (R Core Team 2020) and used package *pls v. 2.7.1* (Mevik et al. 2019) for PLSR modeling.

In our main set of models, we kept chemical traits on a mass basis for consistency with the usual basis on which such traits (except pigments) are measured and reported. However, some traits are most often distributed proportionally to area, and some users may have reasons to prefer area-based estimates (Kattenborn et al. 2019), so have we also made area-based models available (see Table S1 for performance summary statistics). All models (mass- and area-based) are available to download (see *Data availability*).

### External validation

To test how well our PLSR models would transfer to a fully independent dataset, we applied the ensemble of pressed-leaf models for five traits (LMA, LDMC, EWT, N, and C) to pressed-leaf spectra from Cedar Creek, then compared the model-derived trait estimates to measured values. Like most of the CABO dataset, the Cedar Creek dataset comprises trees and herbs from northeastern temperate North America, but it includes an entire functional group (needleleaf conifers) absent among the CABO projects in this study. We collected the spectra with the same instrument and foreoptic as the pressed-leaf spectra in the CABO dataset, but used different sampling, preparation, and measurement protocols. We aimed to see whether the inclusion of a new functional group and the various subtle differences in protocols would affect the models’ performance. Full details on this dataset are provided in Supplementary Materials.

### PLS-DA modeling for species classification

We tested the potential to classify species with fresh-, pressed-, and ground-leaf spectra using partial least-squares discriminant analysis (PLS-DA; Barker & Rayens 2003). We took spectra from the ten most common species in our dataset—all of which are deciduous trees except *A. flexuosa*, which is evergreen. Each was represented by at least 20 specimens (∼480 total). For each tissue type, we divided the full dataset into 60% calibration and 40% validation subsets, stratified by species. In the *R* library *caret v*. *6.0.84* (Kuhn 2020), we trained models on the calibration subset using 10-fold cross-validation repeated 10 times. We chose the number of PLS components during cross-validation by maximizing Cohen’s kappa (κ), which describes the agreement between the true and predicted species identities while accounting for the probability of agreement by chance. Imbalanced training data can bias classification algorithms (Sun et al. 2009), so we used a two-step procedure to balance classes while maintaining enough training data and avoiding overfitting. First, we downsampled within better-represented species classes at random so that all classes had equal size, then chose the number of components (*n*) that maximized κ. Second, we upsampled from less-represented classes at random with replacement so that classes had equal size—again maximizing κ, but restricting the range of components evaluated to no more than the *n* chosen during the downsampling step. We applied the cross-validated PLS-DA model from the upsampling step to the validation subset and summarized its performance using raw classification accuracy and κ.

## Results

### Patterns in traits and reflectance spectra

We saw large variation among samples in each of our target traits within the CABO data, ranging from 1.4-fold variation in C to more than 20-fold variation in traits like lignin, P, K, and Zn (Table 2). The ranges of most traits in our dataset covered a large portion of the global distributions in the TRY dataset, but tended to be narrower at both extremes (Kattge et al. 2020). Many traits—including LMA, LDMC, EWT, cellulose, and many elements—had distributions with a pronounced skew (most often positive). Broadleaf trees tended to have higher LDMC, C, and lignin than other growth forms. Among the herbs, grasses had very high hemicellulose and cellulose and low lignin content, while forbs often had high N. Some of the trait variation was driven by specific projects; for example, *A. flexuosa* in the Warren project tended to have particularly high LMA, Na, and C and low N.

**Table 2:**
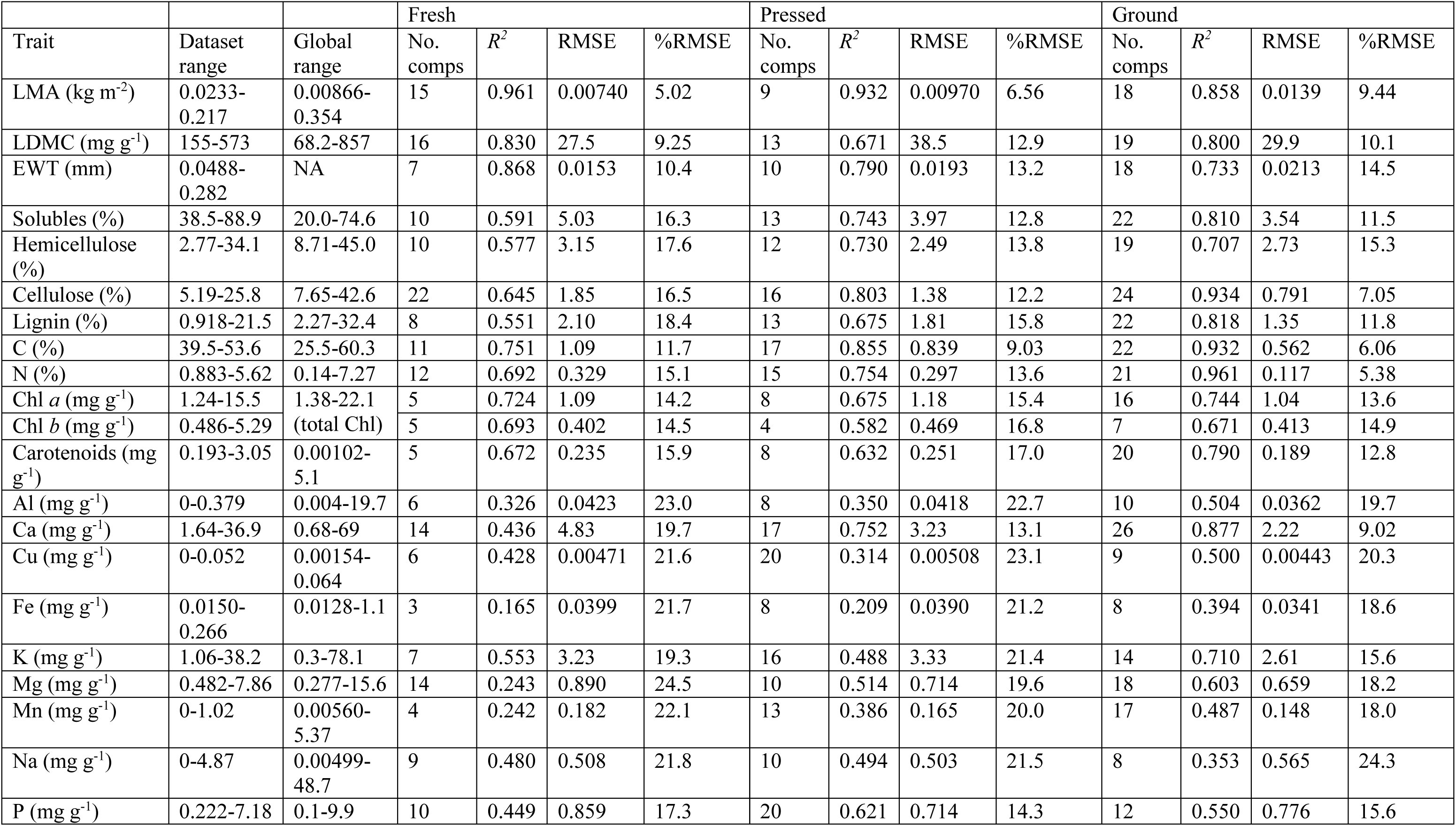
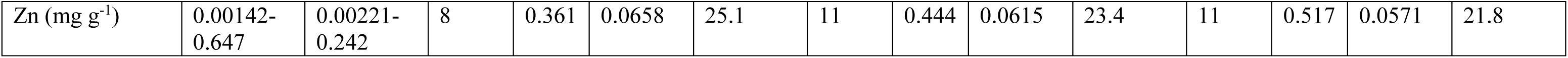
Summary statistics of PLSR model internal validations from full-range fresh-, pressed-, and ground-leaf models. %RMSE is calculated as RMSE divided by the 2.5% trimmed range of measured values within the validation subset. Global ranges are derived from the TRY database (Kattge et al. 2020; accessed 13 April 2021), omitting data with error risk greater than 3. Carbon fractions (solubles, hemicellulose, cellulose, and lignin) and carotenoids are each represented by <1250 points in TRY, which may limit the estimates of the global range. EWT is not available on TRY.

Both pressed and ground leaves had higher median reflectance across nearly the entire spectrum (Fig. 1), as expected based on changes in water content and structure (Carter 1991). Indeed, water absorption features (the largest of which are centered around 1450 and 1930 nm) largely disappeared in pressed and ground leaves. The red edge between the visible and NIR regions was also blunted by both pressing and grinding, causing the global maximum of median reflectance to shift from 872 nm (fresh) to 954 nm (pressed) to 1313 nm (ground).

**Fig. 1:**
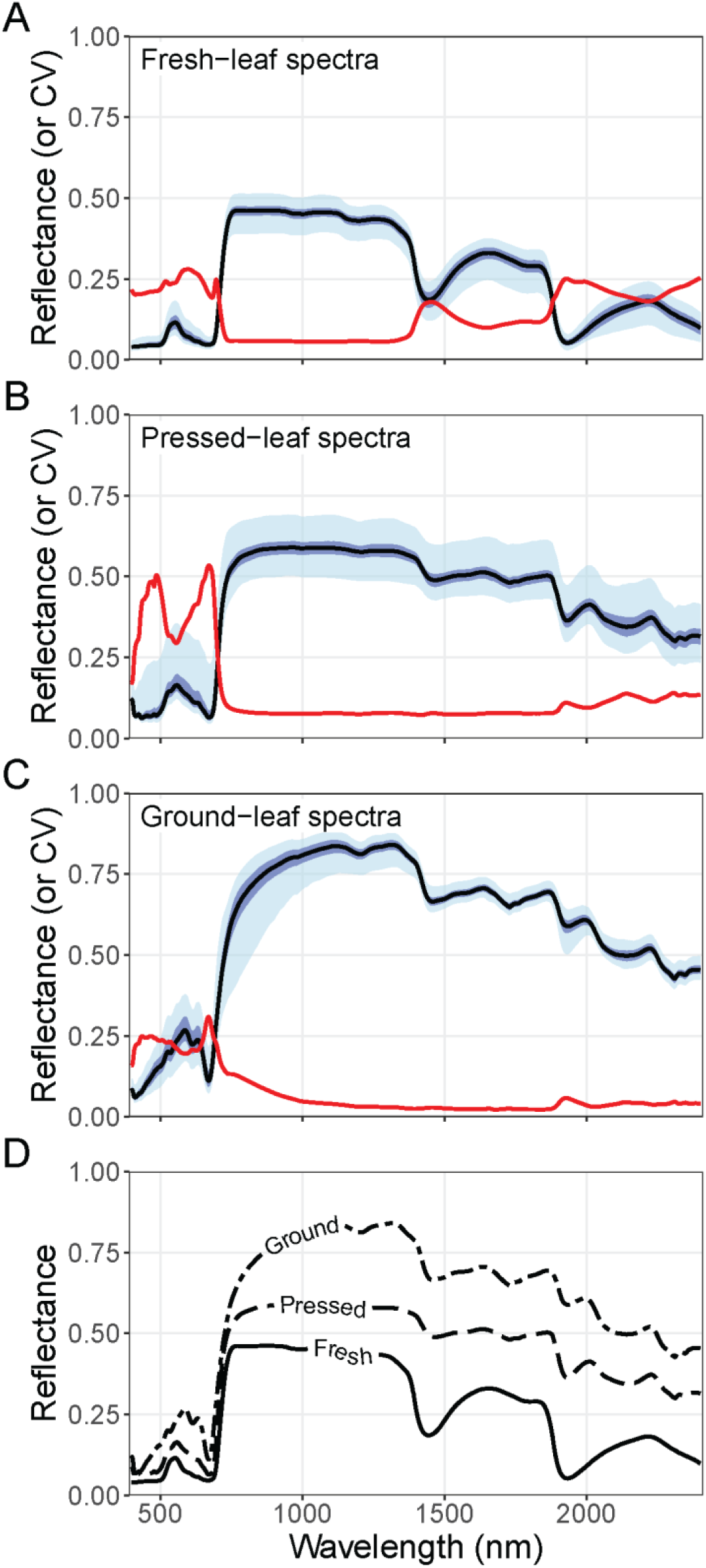
Distributions of spectral reflectance and its coefficient of variation (CV) among fresh (A), pressed (B), and ground (C) leaf samples. A solid black line connects the median reflectance at each wavelength. Dark blue and light blue ribbons denote the middle 50% and 95%. The solid red line shows the coefficient of variation at each wavelength. In (D), the medians of all three are shown for comparison.

Within each tissue type, the coefficient of variation (CV) of reflectance was generally highest where average reflectance was lowest. Across tissue types, pressed-leaf spectra tended to show greater absolute variation in reflectance throughout much of the spectrum, particularly towards the tails of the distribution (e.g. the middle 95 percent in Fig. 1B). The species that have the most exceptionally reflective pressed leaves across the spectrum (mainly *Phragmites australis* (Cav.) Trin. ex Steud., *Phalaris arundinacea* L., and *Asclepias syriaca* L.) do not have particularly reflective fresh leaves, leaving it uncertain why their pressed leaves are so reflective. In contrast, discolored leaves tended to have lower reflectance throughout the visible and NIR ranges (Fig. S5). Unlike pressed leaves, ground leaves showed very low absolute variation in reflectance throughout the SWIR, likely because grinding eliminates variation in leaf structure. However, they showed high variation from 700 to 1100 nm, which may also result from varying degrees of discoloration.

### PLSR modeling for trait estimation

Tested on the internal validation dataset, pressed-leaf models performed best at predicting LMA (*R^2^* = 0.932; %RMSE = 6.56), C (*R^2^* = 0.855; %RMSE = 9.03), and cellulose (*R^2^* = 0.803; %RMSE = 12.2). These traits were followed by a mixture of water-related traits, carbon fractions, and pigments, as well as N and Ca (all *R^2^* = 0.582-0.790; %RMSE = 12.8-17.0). Although some other elements (Mg, P) could be estimated with *R^2^* > 0.5 and %RMSE < 20, most models for other elements showed lower accuracy (Table 2; Figs. 2-3 and S6-12). In most cases, restricted-range (1300-2400 nm) models had similar predictive accuracy to full-range models (Table S2).

**Fig. 2:**
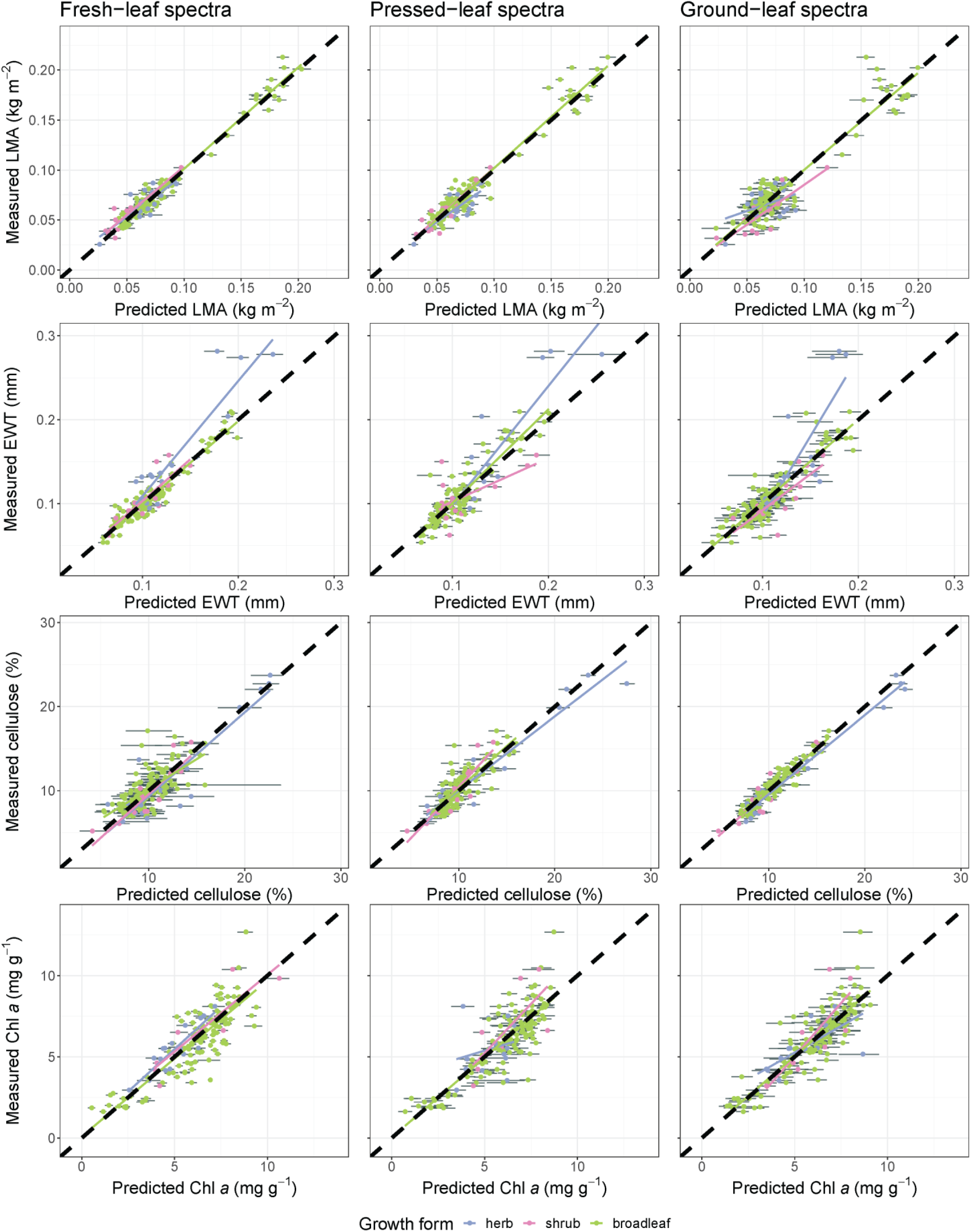
Internal validation results for LMA, EWT, cellulose, and chlorophyll *a* predicted from fresh- (left), pressed- (middle), and ground-leaf (right) spectra. In each panel, each functional group has a separate ordinary least-squares regression line overlaid on top of the thick dashed 1:1 line. The error bars for each data point are 95% confidence intervals calculated from the distribution of predictions based on the ensemble of 100 models produced from jackknife analyses. Figs. 2 and 3 display a subset of traits that represent various functions and span the range of model performance. Plots for remaining traits are in the Supplementary Materials.

**Fig. 3:**
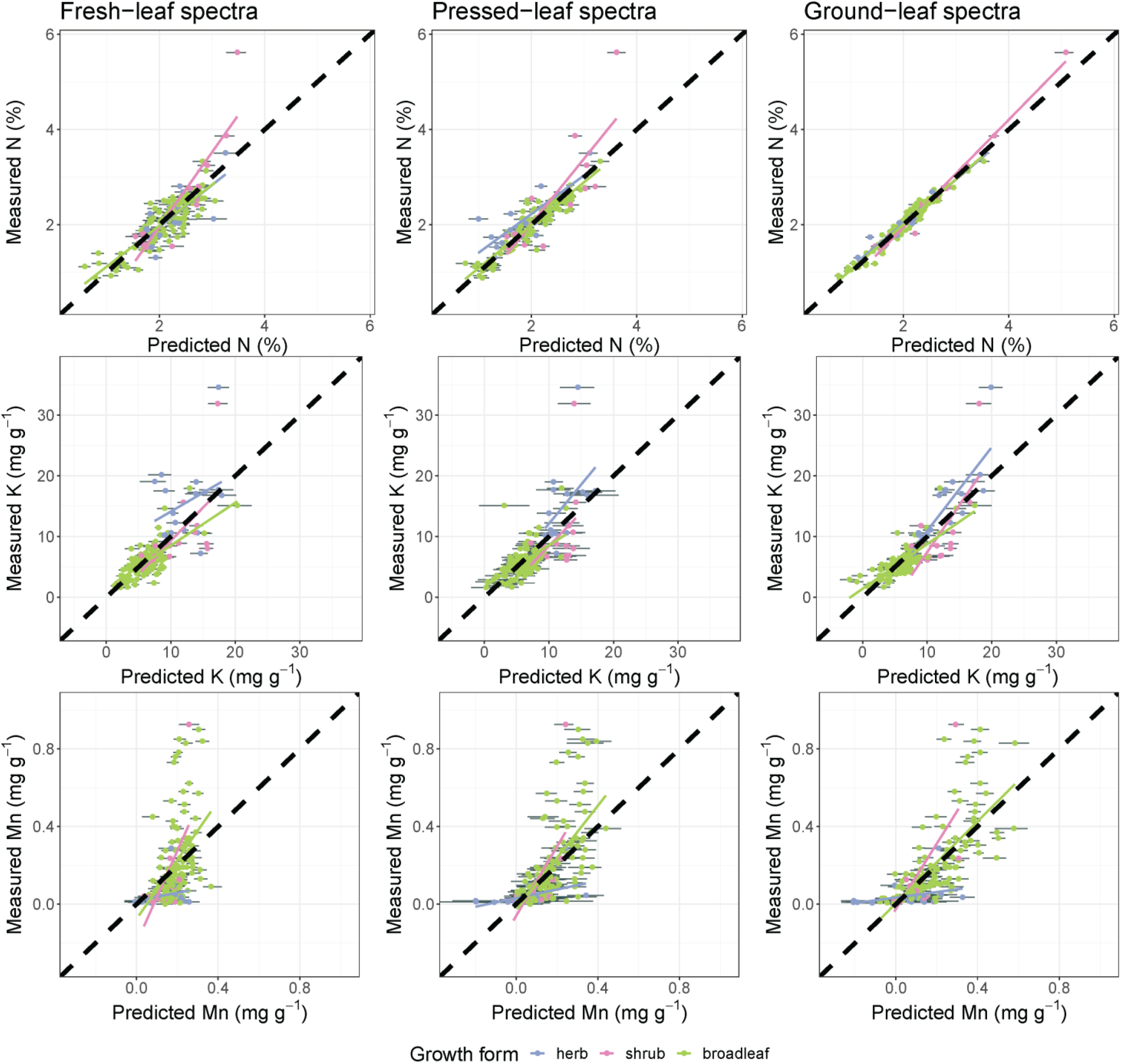
Internal validation results for three elements (N, K, and Mn) predicted from fresh- (left), pressed- (middle), and ground-leaf (right) spectra, displayed as in Fig. 2.

We compared pressed-leaf models to both fresh-leaf and ground-leaf models. The optimal number of components selected to predict each trait was between 3 and 26. For any given trait, ground-leaf models usually had the most components, often followed by pressed-leaf models (Table 2). Fresh-leaf models were best for predicting the structural and water-related traits—LMA, LDMC, and EWT (Figs. 2-3 and S6-12). Ground-leaf models were best for predicting chemical traits, like carbon fractions and most elements. For most traits, pressed-leaf models had intermediate performance, although for some (e.g. pigments, LDMC) both fresh- and ground-leaf models performed better. Statistics from jackknife analyses showed that model performance was more variable for traits that were predicted less accurately (Figs. S13-15). There was no correlation between the magnitude of residuals from pressed-leaf models and our discoloration index for any trait (*p* > 0.05).

For all traits except LMA and Fe, the VIP metric for fresh-leaf spectra showed a global maximum between 710 and 720 nm (Fig. 4 and S16-18)—wavelengths slightly longer than the typical inflection point of the red edge (Richardson et al. 2002). Many traits also show high VIP across the green hump at ∼530-570 nm. Bands in the NIR range were less important for predicting most traits than much of the visible range. The SWIR range was generally important for predicting LMA, EWT, Na, and pigments, and many other traits showed several local peaks of importance, most prominently at about 1880 nm, but also near 1480 and 1720 nm.

**Fig. 4:**
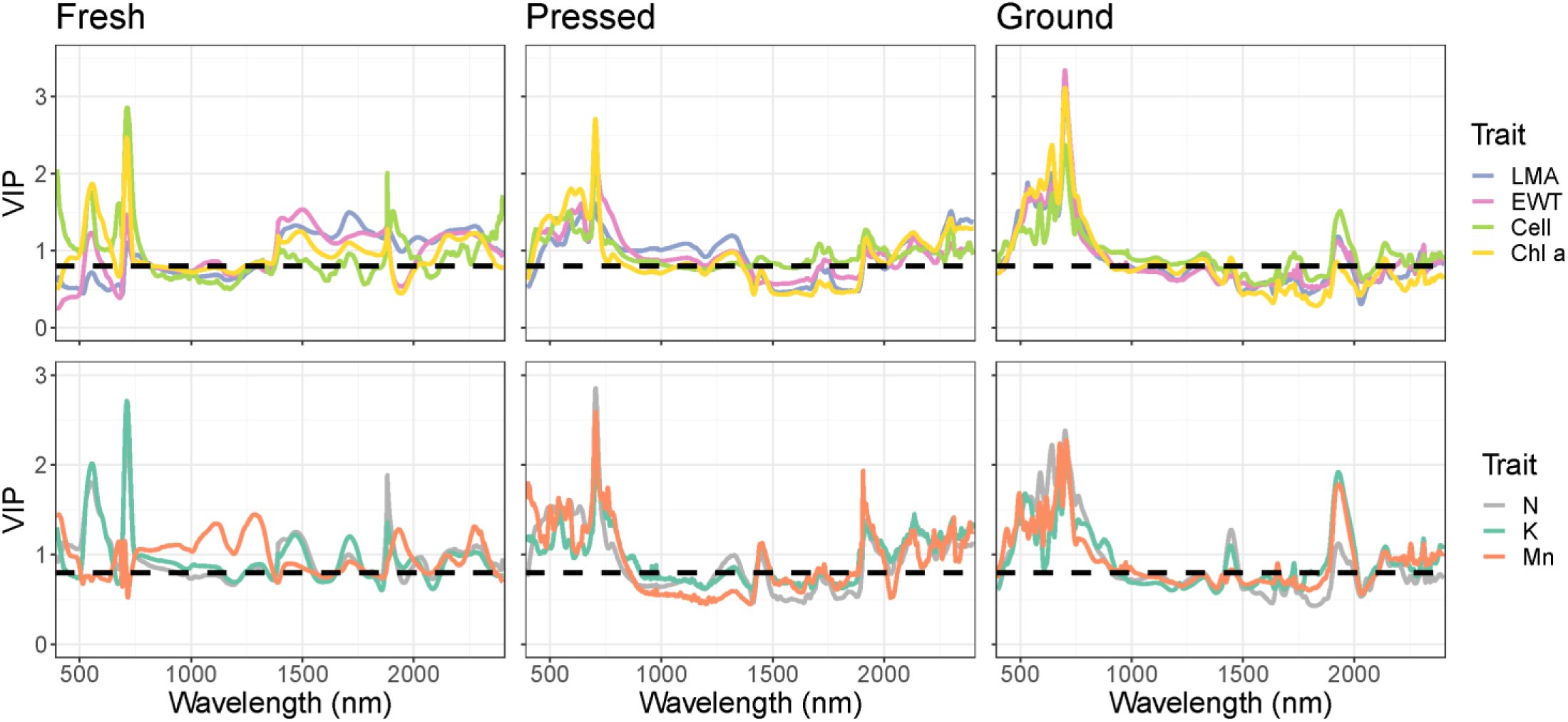
The variable importance of projection (VIP) metric calculated based on fresh- (left), pressed- (middle), and ground-leaf (right) models. As in Figs. 2 and 3, we selected seven traits that represent a range of patterns in VIP across the tissue types (cell = cellulose). The dashed horizontal line at 0.8 represents a heuristic threshold for importance suggested by Burnett et al. (2021). VIP plots for remaining traits are in the Supplementary Materials.

For predicting traits from pressed-leaf spectra, the general trend held that visible reflectance and certain ranges in the SWIR were important for predicting most traits, while the NIR and much of the shorter SWIR (800-1750 nm) were less important (Fig. 4 and S16-18). The red edge peak of importance for most traits was near 705 nm. Other prominent local maxima for many traits lay close to 1440, 1720, 1920, 2130, and 2300 nm. We saw broadly similar patterns in ground-leaf spectra, except that VIP for most traits was lower at longer SWIR wavelengths (2000-2400 nm).

### External validation

For most traits, pressed-leaf model performance on the external validation dataset from Cedar Creek was not quite as strong as the internal validation (Table 3; Fig. 5). For C, the models performed very poorly (*R^2^* < 0.05). Among the remaining traits, *R^2^* ranged from 0.345 (LDMC) to 0.876 (LMA). For N in particular, %RMSE was high (37.6%) due to bias—N concentrations were underestimated for conifers but slightly overestimated for remaining samples.

**Fig. 5:**
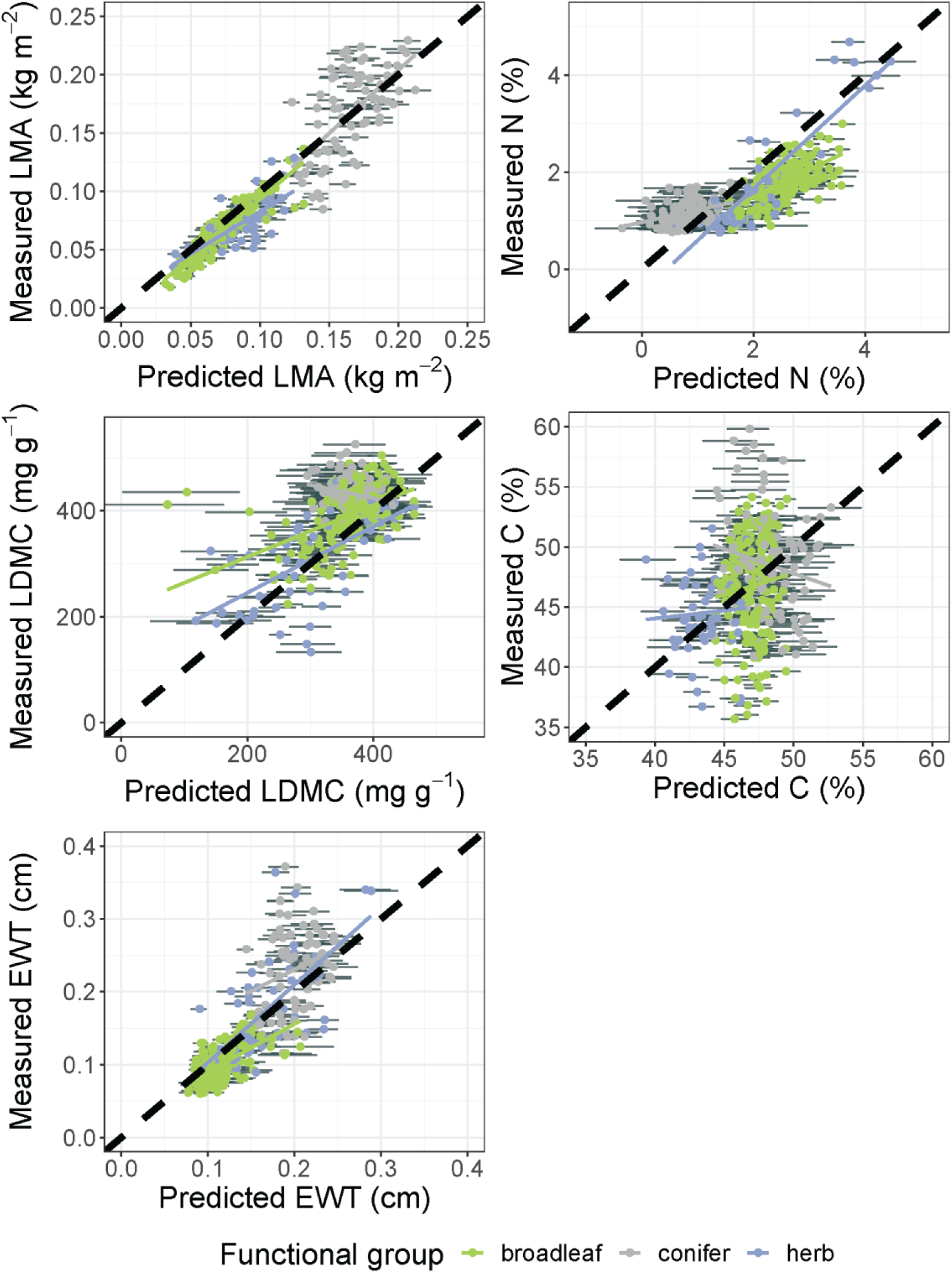
External validation results for various traits in Cedar Creek data predicted using models trained on CABO data. Panels are displayed as in Figs. 2 and 3.

**Table 3:**
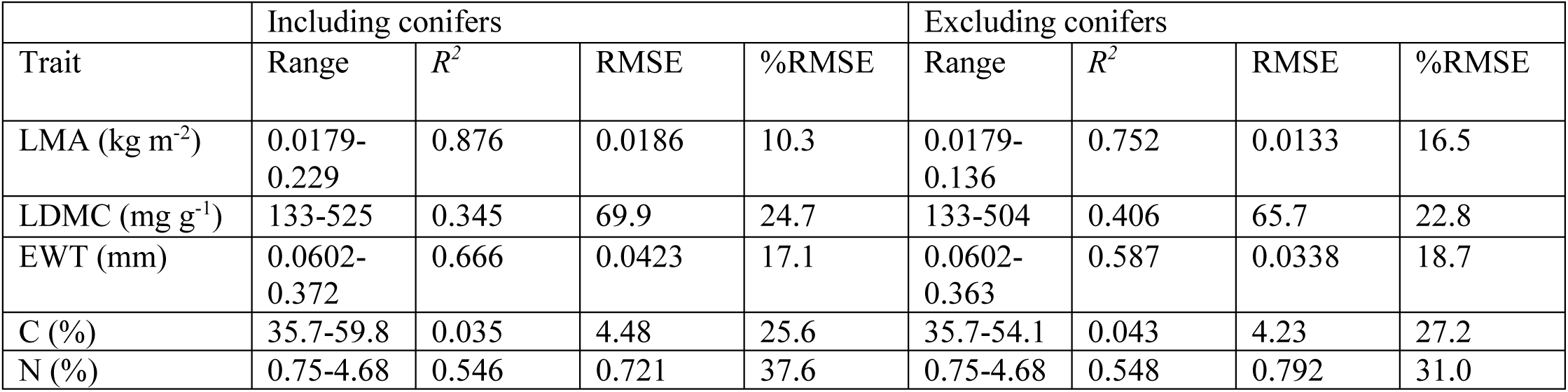
Summary statistics for external validation of pressed-leaf PLSR models. The models were trained on CABO data (see Table 2) and applied to a dataset collected at Cedar Creek.

| Trait | Including conifers |  |  |  | Excluding conifers |  |  |  |
| --- | --- | --- | --- | --- | --- | --- | --- | --- |
|  | Range | *R*<sup>2</sup> | RMSE | %RMSE | Range | *R*<sup>2</sup> | RMSE | %RMSE |
| LMA (kg m<sup>-2</sup>) | 0.0179-0.229 | 0.876 | 0.0186 | 10.3 | 0.0179-0.136 | 0.752 | 0.0133 | 16.5 |
| LDMC (mg g<sup>-1</sup>) | 133-525 | 0.345 | 69.9 | 24.7 | 133-504 | 0.406 | 65.7 | 22.8 |
| EWT (mm) | 0.0602-0.372 | 0.666 | 0.0423 | 17.1 | 0.0602-0.363 | 0.587 | 0.0338 | 18.7 |
| C (%) | 35.7-59.8 | 0.035 | 4.48 | 25.6 | 35.7-54.1 | 0.043 | 4.23 | 27.2 |
| N (%) | 0.75-4.68 | 0.546 | 0.721 | 37.6 | 0.75-4.68 | 0.548 | 0.792 | 31.0 |

Since conifers were absent from the CABO training dataset, we considered whether the models we built would extend to this new functional group. For LDMC, models performed better when excluding conifers (*R^2^* = 0.406, %RMSE = 22.8) than when retaining them (*R^2^* = 0.345; %RMSE = 24.7). In contrast, for LMA and EWT, models performed better when retaining conifers. For LMA in particular, estimates for conifers were quite good, and their extension of the trait range raised *R^2^* (from 0.752 to 0.876) and reduced %RMSE (from 16.5 to 10.3). Restricted-range models yielded somewhat better external validation *R^2^* for N and LDMC both including and excluding conifers (Table S3; Fig. S19). For LDMC in particular, this improvement resulted from improved estimates for extremely discolored samples of *Populus tremuloides* Michx.

### PLS-DA modeling for species classification

PLS-DA models using pressed- and ground-leaf spectra showed near-perfect performance at classifying species (Fig. 6). Models using fresh-leaf spectra were slightly worse but still showed strong performance. The optimal fresh-leaf model, which had 28 PLS components, correctly predicted the taxonomic identity of 175 of the 188 samples in the training dataset (κ = 0.920; *p* < 0.0001). The best pressed-leaf model, which had 37 PLS components, correctly predicted 184 of the 188 samples (κ = 0.975; *p* < 0.0001). The best ground-leaf model, which had 48 PLS components, correctly predicted all 189 samples (κ = 1; *p* < 0.0001). The majority (>70%) of misclassifications were between congenerics.

**Fig. 6:**
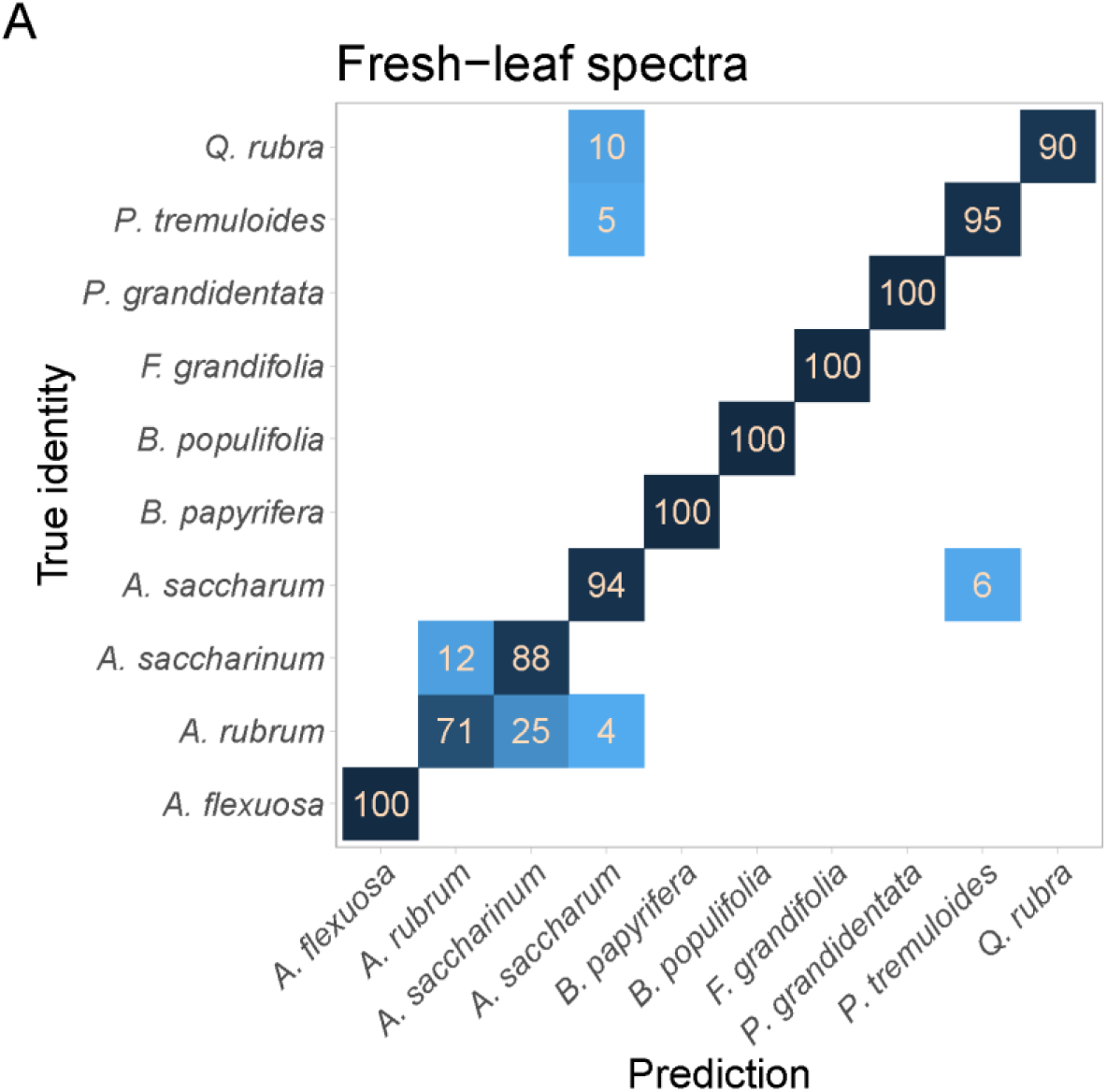
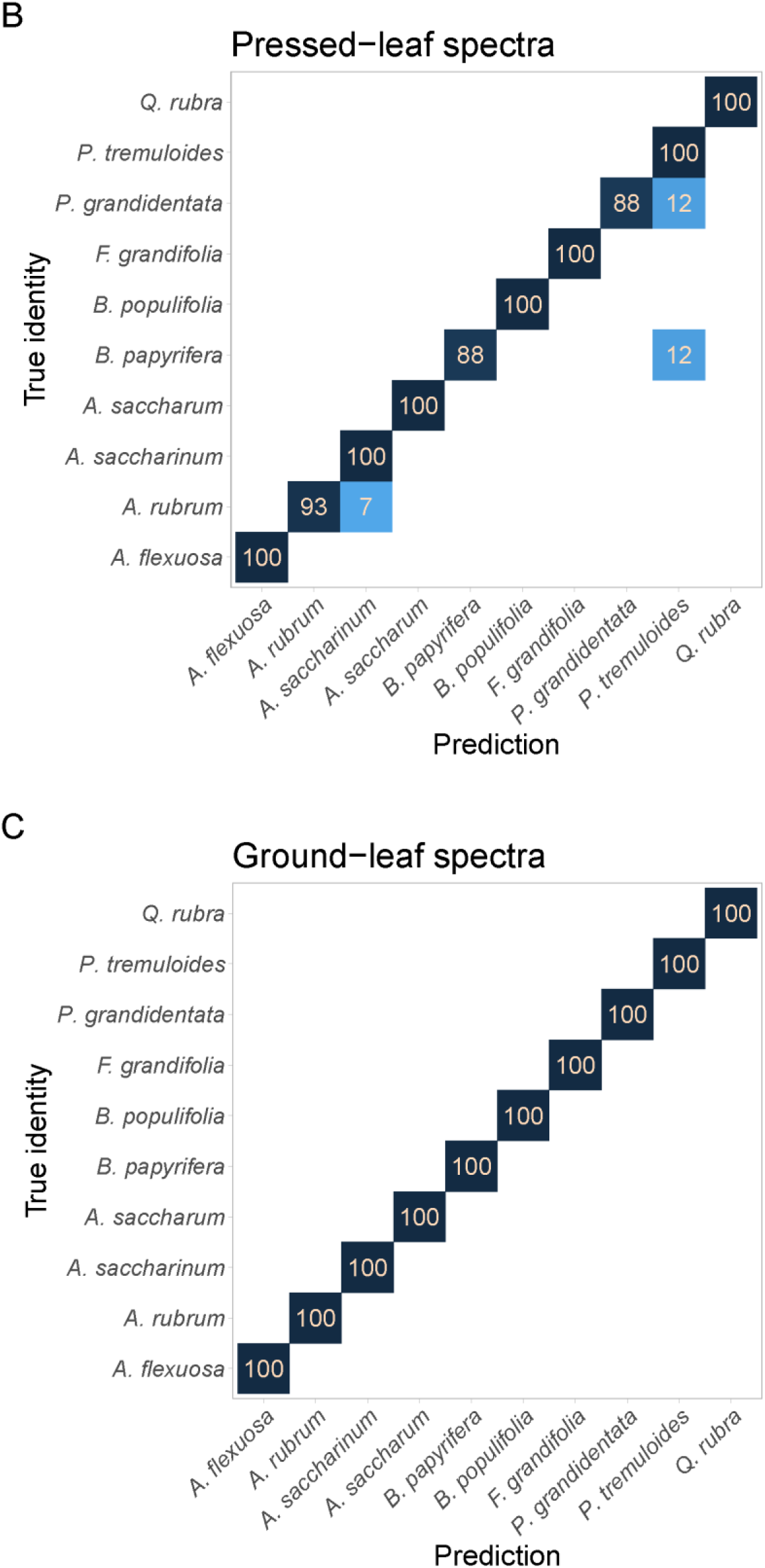
Confusion matrices for partial least-squares discriminant analysis from (A) fresh-, (B) pressed-, and (C) ground-leaf spectra. Rows specify the true species identity, while columns specify the models’ predictions. Each row sums to 100: numbers on the diagonal represent the percentage of specimens of each species that were correctly classified, while off-diagonals represent the percentage misclassified as other species. Full binomials: *Agonis flexuosa*, *Acer rubrum* L., *Acer saccharinum* L., *Acer saccharum* Marshall, *Betula papyrifera* Marshall, *Betula populifolia* Marshall, *Fagus grandifolia* Ehrh., *Populus grandidentata* Michx., *Populus tremuloides* Michx., and *Quercus rubra* L.

## Discussion

We show that we can estimate a wide range of leaf functional traits for 68 woody and herbaceous species from reflectance spectra of pressed leaves (Table 2; Figs. 2-3 and S6-12). Model performance was highest for LMA, C, and N, followed by a mixture of water-related traits, carbon fractions, pigments, and a few important elements (Ca, Mg, and P). Other elements could only be estimated with fairly low accuracy. These results show that pressed-leaf spectra provide an integrative measure of leaf phenotypes, much like fresh-leaf spectra (Cavender-Bares et al. 2017), but with stronger potential to characterize variation in chemical traits. Perhaps as a result, we could use pressed-leaf spectra to classify species as accurately as ground-leaf spectra, and better than fresh-leaf spectra. Our results underscore the potential that using reflectance spectroscopy on herbarium specimens could yield rapid and non-destructive estimates of many functional traits, enabling more expansive studies of trait variation across space and time.

### Comparing PLSR model performance

We compared pressed-leaf models to fresh-leaf and ground-leaf models from the same samples. Our findings about which kind of tissue was best for predicting each trait mostly supported our hypotheses. Ground-leaf spectra showed the strongest performance for most chemical traits, likely because grinding homogenizes the lamina and removes the potentially confounding influence of leaf structure (Table 2). Pressed-leaf spectra showed intermediate performance for most chemical traits, perhaps because, like ground leaves, they lack the major water absorption features that mask the smaller features of other compounds in the SWIR range (Peterson et al. 1988). Contrary to our predictions, ground-leaf spectra performed about as well as fresh-leaf spectra, and better than pressed-leaf spectra, for estimating pigment concentrations. The same factors that provided an advantage to ground-leaf spectra in estimating chemical traits also perhaps explain why pressed- and (especially for LMA) ground-leaf spectra performed worse for estimating water-related and structural traits (LMA, LDMC, and EWT). Pressed-leaf models may represent a good compromise in allowing many traits to be estimated with mostly intermediate but nonetheless quite high accuracy.

Our pressed-leaf models often performed as well as fresh- and ground-leaf models published here and elsewhere. For example, our models for LMA had an RMSE (0.00970 kg m^-2^) lower than many fresh-leaf models from the literature, including Serbin et al. (2019; 0.015 kg m^-2^), Nakaji et al. (2019; 0.015 kg m^-2^), and Streher et al. (2020; 0.051 kg m^-2^). The ground-leaf models in Serbin et al. (2014) had a validation RMSE of 1.4 and 2.4 for cellulose and lignin percentages, comparable to 1.38 and 1.81 for our pressed-leaf models (Table 2). On the other hand, Serbin et al. (2014)’s ground-leaf models for N performed better than our pressed-leaf models (2014; validation RMSE = 0.13 vs 0.297). Our models’ error could be within acceptable bounds for addressing many questions about large-scale ecological or evolutionary patterns that encompass a wide range of trait variation. For some traits (e.g. EWT, N, K, Mn) many of the samples with the greatest errors were at the poorly sampled tails of the measured trait distribution, which suggests that more thorough sampling may be needed to ensure models can make reliable predictions at these extremes (Figs. 2 and 3). Nevertheless, our external validation analyses indicate that our models for some important traits—like LMA, LDMC, EWT, and N, but not C—can transfer reliably to other datasets, and even sometimes to new functional groups (Fig. 5).

### Interpreting PLSR model performance

It may seem perplexing that we could succeed at all in predicting LMA from ground-leaf spectra, or LDMC and EWT from pressed- and ground-leaf spectra. The ability to estimate these traits must not result from the optical expression of the traits themselves. We suggest that we instead sense these traits via their correlations with other traits that have a stronger optical expression. This kind of effect—a “constellation effect” (*sensu* Chadwick & Asner 2016; Nunes et al. 2017)—has been invoked to explain the ability to estimate traits like rock-derived nutrients (Nunes et al. 2017) and δ^15^N (Serbin et al. 2014) that are not known to have strong absorption features in the measured range of wavelengths. However, models that rely on such constellation effects could break down when patterns of trait covariance vary (Kothari & Schweiger 2022), which could make models for certain traits fail beyond the domain of the training data.

The potential role of constellation effects makes it important to interpret how our models work. The VIP metric helps us understand what features are most important, but interpreting it can be challenging because reflectance at a given band is never driven by a single trait. Fresh-leaf VIP for most traits had peaks in the visible and SWIR ranges, with a global maximum at 705 nm (Fig. 4 and S16-18). The pattern of high VIP along the green hump and red edge is common in PLSR models from fresh leaves (Yang et al. 2016; Ely et al. 2019; Streher et al. 2020; Yan et al. 2021). The red edge (particularly 700-725 nm) may be so important because of its sensitivity to both chlorophyll content and leaf structure (Richardson et al. 2002). Much of the visible range was proportionally even more important for pressed- and ground-leaf models, and most of the NIR was less important (except for LMA; Fig. 4 and S16-18). There were multiple small VIP peaks in the SWIR. Although some (e.g. at 1440 and 1920 nm) lie within major water absorption features, any causal link to the leaf’s fresh water content is unlikely for pressed and ground leaves. Many of these peaks also lie near broad absorption features for many components of dry matter, including protein, cellulose, lignin, and starch, which complicates their interpretation (Curran et al. 1989; Fourty et al. 1996).

With some exceptions, the VIP metric showed that the same bands are often important for predicting different traits. This pattern might be taken as an artifact of trait covariance: For example, the three pigment pools covaried strongly (*R^2^* = 0.827-0.969) and had nearly identical VIP across the spectrum (Figs. S16-18). One might take similarities in VIP further to imply that there are a small number of traits whose tight coordination with others underlies the performance of all models through constellation effects. Nevertheless, across the whole dataset, many traits covaried only weakly but still shared VIP patterns. For example, EWT, cellulose, N, and K were not tightly coordinated (*R^2^* = 0.003-0.152) but shared similar patterns of pressed-leaf VIP across the spectrum (Fig. 4), including peaks at 705 and 1920 nm. While VIP is a useful heuristic, it does not show the direction in which a band’s reflectance alters trait estimates; the same bands may matter for different traits in different ways. Here, similarities in VIP do not appear to result solely from strong networks of trait covariance. Nevertheless, the fact that we can estimate traits like LDMC and EWT from pressed-leaf spectra appears to imply some role for trait covariance, perhaps in a more diffuse way.

### PLS-DA modeling for species classification

PLS-DA models showed that fresh-, pressed-, and ground-leaf spectra alike could be used to classify species with perfect accuracy for ground leaves, near-perfect accuracy (>97%) for pressed leaves, and excellent accuracy (>93%) for fresh leaves (Fig. 6). In contrast to prior work that deliberately selected many congenerics (Lang et al. 2015), our most common species were often distantly related. Among the misidentified samples, most were mistaken for congenerics, which implies that related species are more spectrally similar (Schweiger et al. 2018; Meireles et al 2020a). However, past studies using dry leaves have shown great success with closely related species (Lang et al. 2015; Prata et al. 2018) or even populations (Stasinski et al. 2021).

Our analysis reinforces that that pressed- or ground-leaf models might be particularly suited to the task of classifying or delimiting species (Fig. 6). This finding is notable because measuring spectra of pressed leaves in an herbarium is also much simpler than measuring spectra of fresh leaves through an intensive field campaign across the range of a clade. We conjecture that these models have an advantage because drying reveals the absorption features of multiple compounds in the SWIR range that might together allow finer discrimination of species than water content does. Indeed, ground-leaf spectra have greater intrinsic dimensionality than fresh-leaf spectra (Kothari & Schweiger 2022), which suggests they have more independent axes of variation along which species may separate. Our results support the growing practice of using spectra of pressed herbarium specimens in species delimitation and identification (Prata et al. 2018; Draper et al. 2020). However, classification models from pressed or ground leaves have less relevance for research using remotely sensed imagery, which is dominated by fresh leaves.

### The future of spectroscopic trait estimation

Although trait predictions from spectral models are not perfect, they have a few advantages over conventional trait measurements: they (1) can be non-destructive; (2) are fast and require relatively little training; and (3) have very low marginal cost, despite the high capital cost of buying a spectrometer (Costa et al. 2018). These advantages could make it easier to address questions that require large datasets of functional traits. But researchers may be deterred if they must each build their own models tailored to particular uses—and for herbarium specimens, it might not be possible to do the destructive trait measurements often needed to train the models. Ideally, spectral models would be general enough that researchers could confidently use them without further validation, but this aim is not easy to achieve: for several reasons, a model trained on any particular spectral dataset may make poor trait predictions on new data. As with any other technique, the goal for spectroscopic trait estimation is to improve model accuracy and generality as much as they can be jointly improved. Below, we discuss some challenges one by one, particularly as they concern pressed leaves.

One concern is that the new data could be outside the range of traits or optical properties in the training dataset (Schweiger 2020). A general model, if such a thing is possible, would need to represent the vast range of leaf functional traits and optical properties. Another kind of concern about model generality concerns sample preparation before spectral measurements. For example, particle size influences ground-leaf spectra (Foley et al. 1998). For pressed leaves, it may be particularly important to prepare samples in consistent ways that preserve the leaves’ anatomical integrity. In our external validation analyses, we found that pressed-leaf models yielded reasonably accurate predictions of most traits, even though the validation dataset differed in sample preparation protocols and included conifers, which were absent from the training dataset. Nevertheless, even setting aside conifers, external validation for one trait (C) was very poor, and for another (N) showed noticeable bias—enough that researchers might need to develop their own correction factors to use the model in practice.

Another class of challenges concerns spectrometers and their foreoptics. Spectra of fresh leaves can be measured with different foreoptics, including integrating spheres, contact probes, or leaf clips. We used a leaf clip with pressed specimens because mounting delicate pressed leaves in an integrating sphere could damage them. While leaf clips and contact probes often have a higher signal-to-noise ratio, they are less likely to produce consistent measurements among instruments or replicate samples due to variation in viewing geometry and anisotropic surface reflectance (Petibon et al. 2021). The logistical constraint of having to use them on pressed leaves could thus make it harder to compare data among instruments. In theory, the greater inconsistency of leaf clip measurements could have reduced the performance of our pressed-leaf models compared to our integrating sphere-based fresh-leaf models, but we still found that the former performed better for most chemical traits.

Another challenge is that while many herbarium specimens are glued to a paper backing, measuring reflectance with a leaf clip or probe usually requires placing a black absorbing background under the sample to keep transmitted light from being reflected back into the sensor. When unattached leaves are not available, using spectra from these specimens may require new methods to correct for reflectance from the mounting paper.

Spectrometers and their software also vary subtly in their sensors and techniques for processing spectra (Castro-Esau et al. 2006), and in some cases researchers must take steps to reconcile spectra measured from different instruments (Meireles et al. 2020b). Assuming that these kinds of technical challenges can be overcome, our results and others encourage confidence about building general models to estimate traits from a wide variety of plants (Serbin et al. 2019). The creation of open libraries for spectral data (like EcoSIS, https://ecosis.org/; or the CABO data portal, https://data.caboscience.org/leaf) and spectral models (like EcoSML; https://ecosml.org/) will contribute to this goal. Lastly, we note that many of the same concerns about discrepancies among sampling and measurement protocols could arise when using existing spectral libraries to aid in species identification (Draper et al. 2020).

### Implications for herbarium-based research

A particular challenge for herbarium specimens is that their optical and chemical properties (especially light-sensitive pigments) may degrade during preparation or storage. Such degradation could make it hard to distinguish changes in the traits of living plants over time from changes in storage. Even in this study, where no specimens were collected before 2017, many underwent visible changes in color, including browning or blackening; ∼12% were scored at 2 or higher, with large variation among species (e.g. 42% of *Populus grandidentata* specimens, but 0% of *Betula papyrifera* specimens). We found little evidence that such discoloration hinders trait estimation: Both the results of our discoloration analyses and the similar performance of full- and restricted-range models suggest that PLSR is flexible enough to predict traits despite the variable influences of discoloration in our specimens. This capacity likely depends on using samples for model calibration that show a similarly wide range of discoloration.

Our specimens were collected no more than three years before measurement, but ecologists may want to use specimens collected decades ago. While our findings give reason to be optimistic that properly calibrated models could return accurate estimates of many traits from old or discolored specimens, it remains untested whether there are any limits to this potential. In general, not much is known about long-term changes in specimen chemical (aside from DNA) or optical properties (Lang et al. 2018). Color changes are known to accelerate under certain preparation and storage conditions—including exposure to some chemical preservatives and high light, drying temperature, or humidity (Bridson & Forman 1999; Metsger & Byers 1999)—which it may be worth avoiding when possible. Long-term studies of specimens—perhaps subjected to varying preparation and storage techniques—could establish how chemical and optical properties change over time and help refine these guidelines further.

Some of the challenges we describe pertain to projects that would measure spectra on samples already collected, but spectroscopy—like other novel uses for herbarium specimens—could also prompt changes in collection practice. For example, it underscores the potential value of gathering and storing extra leaf material (e.g. in fragment packets), which would circumvent the challenge of measuring mounted leaves and aid destructive analyses of herbarium specimens (Heberling 2021). We propose that herbaria could also incorporate spectroscopy into their operations by measuring incoming specimens shortly after pressing, which could mitigate the challenges caused by mounting and degradation.

Linking spectral data measured on herbarium specimens to the digital record of the voucher could be a powerful tool to enable data synthesis, but it may require new informatic tools (Draper et al. 2020). The hyperdimensionality of the reflectance spectrum could make it hard to accommodate within existing standards like the Darwin Core (Wieczorek et al. 2012), at least without extensions. One could link records to external spectral databases like EcoSIS or SPECCHIO (Hueni et al. 2020), which are also designed to store metadata about instrumentation and processing. We would advocate for coordination between herbarium managers and researchers who use reflectance spectroscopy, which could build agreement about best practices for spectral measurement and curation and allow data to be synthesized and compared across research groups.

We show that non-destructively measured pressed-leaf spectra retain much of the information about many leaf functional traits found in fresh-leaf spectra. While validating this technique on older specimens will require extensive further research, our findings suggest that reflectance spectroscopy could allow herbaria to take on a greater role in plant functional ecology and evolution. Our study has far-reaching implications for capturing the wide range of functional and phenotypic information in the world’s preserved plant collections.

## Acknowledgements

We performed this research on the ancestral and contemporary land—mostly unceded—of Native people: primarily the Kanien’kehá꞉ka (Mohawk), Omàmiwinini (Algonquin), and Abenaki First Nations for the Beauchamp-Rioux, Dessain, and Boucherville projects, the Noongar peoples for the Warren project, and the Dakota and Ojibwe peoples for the Cedar Creek dataset. For permissions and assistance, we thank the Jardin botanique de Montréal; la Station de biologie des Laurentides (SBL) de l’Université de Montréal; l’Institut de recherche d’Hydro-Québec (IREQ); the National Capital Commission for sampling at Mer Bleue Bog; la Société des établissements de plein air du Québec (SÉPAQ) for sampling at Parc national d’Oka, Parc national du Mont-Saint-Bruno, and Parc national des Îles-des-Boucherville; and the Parks and Wildlife Service of Western Australia for sampling in D’Entrecasteaux National Park. We thank Antoine Mathieu, Alexandra Massey, Florence Blanchard, Elisabeth Hardy-Lachance, Zachary Bélisle, Sandra Jooty, Myriam Cloutier, Fabien Cichonski, Madeleine Trickey-Massé, Vincent Fournier, Aurélie Dessain, Xavier Guilbeault-Mayers, Jocelyne Ayotte, Megan Erding, and especially Sabrina Demers-Thibeault for contributing to trait measurements. Carole Sinou, Phil Townsend, Dudu Meireles, Mason Heberling, and members of the Cavender-Bares lab and the Laboratory of Plant Functional Ecology (LEFO) contributed to insightful discussions. Data from Canadian and Australian sites were collected as part of the Canadian Airborne Biodiversity Observatory (CABO) funded by NSERC Discovery Frontiers grant 509190-2017, as well as NSERC Discovery grants (RGPIN-2014-06106, RGPIN-2019-04537) and a Canada Research Chair to EL. Cedar Creek collections were supported by the NSF/NASA (DEB-1342778) and the NSF ASCEND Biology Integration Institute (DBI-2021898). Cedar Creek is supported by the NSF Long-Term Ecological Research program, currently under DEB-1831944. Travel funding was provided by an International Thesis Research Travel Grant from the University of Minnesota’s Graduate School and a grant distributed through an NSF-funded Research Coordination Network (DEB-1745562). SK was supported by an NSF Graduate Research Fellowship (Grant No. 00039202) and a UMN Doctoral Dissertation Fellowship. RBR was supported by NSERC through the Canada Graduate Scholarships— Master’s program and the FRQNT through a Master’s Research Scholarship.

## Conflict of interest statement

The authors have no conflicts of interest to declare.

## Author contributions

SK, JCB, and EL conceived the ideas and designed the methodology. RBR and SK collected the data. SK analyzed the data and wrote the first draft of the manuscript. All authors made substantial contributions to further drafts and gave final approval for publication.

## Data availability

All fresh-leaf spectral data are available through the CABO data portal (https://data.caboscience.org/leaf). Upon publication, we will upload other spectral data, as well as metadata and trait data, to the Ecological Spectral Information System (EcoSIS, https://ecosis.org/), and upload models to the Ecological Spectral Model Library (EcoSML, https://ecosml.org/). At that stage, we will update this section accordingly.

## Supplementary Materials

### Sampling

For each sample, we collected a large group of sunlit leaves (>3 h estimated sun exposure per day) at about the same vertical position—for trees, the same branch or neighboring branches from the same individual. We avoided leaves with noticeable damage from herbivores or pathogens. We measured fresh-leaf spectra and fresh-leaf mass in the field, froze leaf disks for pigment analyses, and transported samples in a cooler to the lab. We then measured other whole-leaf traits and divided the remainder of the sample into two groups: (1) a portion to be pressed, and (2) a portion whose leaves were dried and ground. We measured pressed-leaf spectra on the former portion prior to mounting, and ground-leaf spectra and chemical traits on the latter. Although these are distinct subsets of leaves, we treat them as identical in their trait values since they are divided at random from the larger, relatively homogeneous bulk sample. A minimum of one representative pressed specimen of each species at each site and date is prepared as a specimen at the Marie-Victorin Herbarium in Montréal, Québec, Canada.

We classified species’ growth forms following the Database of Vascular Plants of Canada (VASCAN; Desmet & Brouillet 2013). We classified *Agonis flexuosa* as a broadleaf tree and manually disambiguated some species listed as adopting either shrub or three growth forms.

### Spectral measurements

For the Beauchamp-Rioux, Boucherville, and Warren projects, we measured fresh-leaf directional-hemispherical reflectance spectra on each sample in 2018 using an HR-1024i spectroradiometer outfitted with a DC-R/T integrating sphere from Spectra Vista Corporation (SVC; Poughkeepsie, NY, USA). We measured spectra on the adaxial surface of six leaves or, for small or narrow leaves, single-layer leaf arrays following the protocol in Noda et al. (2013) as adapted for our instrument (Laliberté & Soffer 2018a; Laliberté and Soffer 2018b). For the Dessain project, we measured fresh-leaf reflectance spectra in 2017 using a FieldSpec 4 spectroradiometer equipped with an RTS-3ZC integrating sphere from Analytical Spectral Devices, Inc. (ASD; Boulder, CO, USA) following the leaf spectroscopy protocol from the Global Airborne Observatory (https://gao.asu.edu/spectranomics-protocols). We collected spectra from a variable number of leaves or arrays per sample—six in most cases, but as few as three or as many as eighteen from certain samples. For all projects, each reflectance measurement was calibrated against a white Spectralon 99% reflectance panel and corrected for stray light.

Each pressed specimen was dried in a plant press between sheets of newspaper in a setting with constant air flow and at most gentle heating (≤ 30 °C) for a minimum of 48 h. For all projects, we measured reflectance spectra on the adaxial surface of pressed, intact leaf samples using a PSR+ 3500 spectroradiometer with a leaf clip foreoptic (Spectral Evolution, Haverhill, MA) between six months and three years after collection. We calibrated reflectance against a white Spectralon reflectance panel and measured 3-7 spectra per sample. Although we pressed samples until dry, some nevertheless dried in a way that created an uneven leaf surface. At the time of measurement, we noted any visible discoloration that samples underwent during drying or storage, including browning or blackening (Figs. S2-5). In general, we aimed to avoid uneven or discolored areas when measuring spectra, although it was sometimes not entirely possible when discoloration was pervasive.

Each ground-leaf bulk sample was oven-dried at 65 °C for three days, ground using a cyclone mill with 2 mm mesh, redried at 65 °C, and stored for chemical analyses. For all projects, we also measured ground-leaf spectra using the PSR+ 3500 spectroradiometer with a benchtop reflectance probe foreoptic that pressed loose powder into a smooth, even pellet. For each sample, we added at least 0.6 g of leaf powder into a sample tray; in preliminary tests, adding additional material did not change the spectra, suggesting that transmittance was close to zero. We calibrated reflectance against a white Spectralon reflectance panel placed in an identical sample tray. We measured three spectra per sample on the same pellet, turning the sample tray 120° between the first two spectra, and loosening and mixing the powder before reshaping the pellet between the second and third spectra.

A small number of specimens did not have usable spectra at all three stages, usually either due to lack of enough pressed or ground material or to poor spectral quality at any stage. To ensure fair comparisons among the stages, we removed any sample from the dataset that was not available for at least two stages. Among the remaining samples, any sample that was not available for all three stages (<4%) was placed in the training subset for PLSR analyses.

Our spectral processing pipeline varied by the type of data. Although the actual spectral resolution was more variable and coarser, the ASD Field Spec 4 and Spectral Evolution PSR+ 3500 spectrometers’ software automatically resampled the spectra to 1 nm resolution and interpolated over the overlap region between sensors (Schweiger & Laliberté 2020). For data from the SVC HR-1024i spectrometer, we performed these steps using linear interpolation. For pressed-leaf spectra only, we detected jumps in reflectance at the sensor overlap region and removed them using the function *match_sensors()* in *R* package *spectrolab v. 0.0.10* (Meireles et al. 2017). Next, we averaged all spectra for a given sample and tissue type (fresh, pressed, or ground). Spectra measured with integrating spheres tend to have greater noise, especially at the ends. To reduce noise in fresh-leaf spectra, we applied a Savitzky-Golay filter using *R* package *signal 0.7.6* (Signal Developers, 2013), with varying order and length: order 3 and length 21 from 350-715 nm, order 3 and length 35 from 715-1390 nm, order 3 and length 75 from 1390-1880 nm, and order 5 and length 175 from 1880-2500 nm. Finally, we trimmed all spectra to 400-2400 nm, removing the extremes where sensors show greater noise. We did all spectral processing in *R v. 3.6.3* (R Core Team 2020) using *spectrolab v. 0.0.10* (Meireles et al. 2017).

### Leaf trait measurements

We performed all leaf trait measurements excluding petioles, but including the rachis for compound leaves, since the rachis is functionally analogous to the midrib of a simple leaf. We measured the following leaf structural and chemical traits: Leaf mass per area (LMA; kg m^-2^), leaf dry matter content (LDMC; mg g^-1^), equivalent water thickness (EWT; mm), carbon fractions (soluble cell contents, hemicellulose, cellulose, and lignin) on a mass basis, pigments (chlorophyll *a*, chlorophyll *b*, and total carotenoids) on a mass basis, and concentrations of a variety of elements (Al, C, Ca, Cu, Fe, K, Mg, Mn, N, Na, P, Zn).

From a subset of the leaves for each sample whose fresh-leaf spectra we measured, we used a 7 mm cork borer to punch leaf disks (or cut fragments of narrow leaves) in the field and stored them on ice in the darkness before transferring them to a -80 °C freezer in the lab. We extracted pigments under darkness in an MgCO_3_-MeOH solution and estimated pigment concentrations using a spectrophotometric protocol on a SPECTROstar Nano microplate reader (BMG LABTECH, Ortenburg, Germany; Girard et al. 2020).

From a separate subset of leaves from each sample, we weighed fresh mass in the field shortly after collection. We then rehydrated the leaves in sealed plastic bags with damp paper towels for at least 12 h and weighed them for rehydrated mass. We scanned leaves and estimated their area using WinFOLIA (Regent Instruments, Québec, QC, CA). Lastly, we dried them for at least 72 h in a drying oven at 65°C before reweighing for dry mass. We calculated LDMC as total dry mass divided by total fresh mass and LMA as total dry mass divided by total rehydrated area (Laliberté 2018). We further calculated EWT from LDMC and LMA. For the Dessain project alone, fresh mass in the field was not available, so we used rehydrated mass to calculate LDMC and EWT, which may lead our reported values to underestimate LDMC and overestimate EWT. These biases are likely to be relatively small since, in the remaining projects, relative water content was generally close to 100% (median: 89.5%, 2.5–97.5^th^ percentile: 79.2–97.8%).

We analyzed ground-leaf bulk samples for N and C using a Vario MICRO Cube combustion analyzer (Elementar, Langenselbold, Germany; Ayotte et al. 2019). We analyzed the same samples for other elements (Al, Ca, Cu, Fe, K, Mg, Mn, Na, P, Zn) using nitric acid digestion followed by inductively coupled plasma-optical emission spectrometry (ICP-OES) using an Optima 7300CV (PerkinElmer, Waltham, MA, USA). We removed five data points from the ICP-OES output as extreme outliers that were not representative of the range of concentrations of their respective elements across our samples. We also set values below the detection limit, which sometimes returned as negative concentrations, to zero.

We also analyzed the ground-leaf samples for carbon fractions using an ANKOM 2000 Fiber Analyzer (Ankom Technology, Macedon, New York, USA) to perform a sequence of digestions (Ayotte & Laliberté 2019). The first digestion in a heated neutral detergent washes off soluble cell contents. The second digestion in a heated acidic detergent removes hemicellulose and bound proteins—which, for simplicity, we just refer to as hemicellulose. The third digestion in 70% sulfuric acid removes cellulose, leaving behind lignin and recalcitrant compounds. We used the change in each bag’s mass between stages to determine the mass of each fraction. We ashed all samples and used ash content as a correction factor to calculate the ash-free lignin fraction.

### External validation dataset

From August to November 2018, we collected 333 samples for an external validation dataset at Cedar Creek Ecosystem Science Reserve (East Bethel, MN, USA). Most were from the Forests and Biodiversity (FAB) experiment (Grossman et al. 2017), the Biodiversity in Willows and Poplars (BiWaP) experiment (Grossman & Cavender-Bares 2019), and the Big Biodiversity (BioDIV or e120) experiment (Tilman et al. 1997), although some were from other parts of the site.

Each sample included multiple leaves from the same individual, taken when possible from a nearby position on the plant. Unlike in the CABO dataset, we collected samples from any part of an individual, not just sunlit leaves. From each sample, we designated one leaf for fresh mass to be measured immediately following collection; this leaf (‘fresh-mass subsample’) was stored separately and its spectra were labeled to keep track of its identity. (For conifers or other plants with small or narrow leaves, we often included multiple leaves in this fresh mass sample.) We took and flash-froze 3-5 hole punches from another leaf (or group of leaves) for pigment analyses (‘pigment subsample’; pigment data not shown); this leaf and its spectra were also labeled separately. We collected more leaves to have enough tissue for a bulk chemical sample, but did not keep track of them individually.

In the field, we measured multiple fresh-leaf spectra per sample (not shown here) using a PSR+ 3500 spectroradiometer with a leaf clip foreoptic (Spectral Evolution, Haverhill, MA) and placed the leaves in sealed plastic bags with a damp paper towel. These bags were kept in coolers until the end of the sampling day. We weighed the fresh-mass subsample immediately upon return to the lab and kept all leaves in a refrigerator until we scanned the fresh-mass and pigment subsamples within the next few days. After scanning, we placed all leaves in flat paper bags and pressed them for several hours in a plant press before removing them and placing them in a drying shed at 40 °C for at least one week. This deviates from most standard herbarium protocols, including the one used in the CABO project, where leaves are kept in the press while drying. After we dried the leaves, we weighed the fresh-mass and pigment subsamples. Certain species, especially *Populus tremuloides*, seemed especially prone to discoloration during sample preparation.

In May 2020, we measured pressed-leaf spectra using a PSR+ 3500 spectroradiometer with a leaf clip foreoptic. We did not use a specific protocol for small or narrow leaves, but aimed to arrange leaves in a way that minimized overlaps or gaps. We removed spectra with especially low (<40%) or high (>80%) peak reflectance, and any with major visible gaps in the sensor overlap regions. We measured three spectra per subsample except in cases where getting high-quality spectra proved especially difficult. We then removed the petioles (when applicable) and put leaves in scintillation vials with ball bearings to grind the tissue in a paint shaker, combining all subsamples from a given sample. We then dried samples again and measured C and N with an elemental analyzer (Costech ECS 4010 Analyzer, Valencia, California, USA).

For estimating LMA, LDMC, and EWT, we only used the spectra from the fresh-mass subsample. For C and N, we used all spectra from a given sample, ignoring the subsample labels. As among the CABO pressed-leaf spectra, we first matched sensors, then averaged spectra within the relevant sample or subsample, and lastly trimmed them to 400-2400 nm.

We applied the ensemble of models resulting from our 100× jackknife analysis on the pressed-leaf CABO data and applied them to the Cedar Creek data. We also applied the model ensemble from the pressed-leaf restricted-range (1300-2500 nm) CABO data to the restricted-range Cedar Creek data. We calculated summary statistics and plotted results as in the internal validation.

## Supplementary tables

**Table S1:** Summary statistics of PLSR model internal validation for area-based chemical traits from full-range fresh- and pressed-leaf models. %RMSE is calculated as RMSE divided by the 2.5% trimmed range of measured values within the validation subset.

| Trait | Fresh |  |  |  | Pressed |  |  |  |
| --- | --- | --- | --- | --- | --- | --- | --- | --- |
| | No. comps | $R^2$ | RMSE ($\times 10^6$) | %RMSE | No. comps | $R^2$ | RMSE ($\times 10^6$) | %RMSE |
| Solubles (g cm<sup>-2</sup>) | 9 | 0.948 | 614 | 5.88 | 8 | 0.915 | 791 | 7.56 |
| Hemicellulose (g cm<sup>-2</sup>) | 17 | 0.688 | 258 | 15.8 | 11 | 0.754 | 228 | 14.0 |
| Cellulose (g cm<sup>-2</sup>) | 21 | 0.858 | 158 | 10.3 | 15 | 0.902 | 133 | 8.58 |
| Lignin (g cm<sup>-2</sup>) | 13 | 0.898 | 167 | 8.17 | 16 | 0.916 | 151 | 7.41 |
| C (g cm<sup>-2</sup>) | 16 | 0.969 | 349 | 4.48 | 10 | 0.942 | 475 | 6.07 |
| N (g cm<sup>-2</sup>) | 19 | 0.753 | 20.7 | 13.9 | 14 | 0.761 | 19.9 | 13.3 |
| Chl *a* (g cm<sup>-2</sup>) | 6 | 0.588 | 6.93 | 17.2 | 7 | 0.570 | 7.00 | 17.3 |
| Chl *b* (g cm<sup>-2</sup>) | 6 | 0.535 | 2.56 | 18.6 | 6 | 0.502 | 2.63 | 19.0 |
| Carotenoids (g cm<sup>-2</sup>) | 9 | 0.562 | 1.36 | 17.6 | 9 | 0.477 | 1.48 | 19.0 |
| Al (g cm<sup>-2</sup>) | 16 | 0.281 | 0.362 | 31.3 | 7 | 0.255 | 0.372 | 31.9 |
| Ca (g cm<sup>-2</sup>) | 15 | 0.529 | 39.6 | 18.3 | 17 | 0.75 | 29.1 | 13.4 |
| Cu (g cm<sup>-2</sup>) | 5 | 0.273 | 0.0302 | 21.6 | 8 | 0.132 | 0.0317 | 31.6 |
| Fe (g cm<sup>-2</sup>) | 2 | 0.040 | 0.265 | 25.3 | 7 | 0.080 | 0.261 | 24.9 |
| K (g cm<sup>-2</sup>) | 9 | 0.524 | 18.1 | 17.2 | 18 | 0.553 | 16.0 | 16.6 |
| Mg (g cm<sup>-2</sup>) | 7 | 0.564 | 6.72 | 16.8 | 11 | 0.707 | 5.50 | 13.7 |
| Mn (g cm<sup>-2</sup>) | 4 | 0.162 | 1.44 | 23.5 | 15 | 0.402 | 1.23 | 20.1 |
| Na (g cm<sup>-2</sup>) | 5 | 0.701 | 6.06 | 15.2 | 11 | 0.767 | 5.46 | 13.7 |
| P (g cm<sup>-2</sup>) | 3 | 0.338 | 8.20 | 24.1 | 13 | 0.553 | 6.59 | 19.2 |
| Zn (g cm<sup>-2</sup>) | 10 | 0.336 | 0.495 | 22.6 | 14 | 0.480 | 0.44 | 20.0 |

**Table S2:** Summary statistics of PLSR model internal validation from pressed-leaf spectra restricted to 1300-2500 nm.

| Trait | No. comps | $R^2$ | RMSE | %RMSE |
| --- | --- | --- | --- | --- |
| LMA (kg m<sup>-2</sup>) | 8 | 0.922 | 0.0105 | 7.08 |
| LDMC (mg g<sup>-1</sup>) | 11 | 0.703 | 36.9 | 12.4 |
| EWT (mm) | 11 | 0.781 | 0.0194 | 13.3 |
| Solubles (%) | 11 | 0.759 | 3.88 | 12.5 |
| Hemicellulose (%) | 10 | 0.725 | 2.51 | 13.9 |
| Cellulose (%) | 10 | 0.783 | 1.44 | 12.6 |
| Lignin (%) | 11 | 0.670 | 1.83 | 15.9 |
| C (%) | 11 | 0.864 | 0.825 | 8.89 |
| N (%) | 11 | 0.741 | 0.303 | 13.9 |
| Chl *a* (mg g<sup>-1</sup>) | 7 | 0.631 | 1.26 | 16.4 |
| Chl *b* (mg g<sup>-1</sup>) | 7 | 0.570 | 0.475 | 17.0 |
| Carotenoids (mg g<sup>-1</sup>) | 7 | 0.652 | 0.244 | 16.5 |
| Al (mg g<sup>-1</sup>) | 6 | 0.231 | 0.0452 | 24.5 |
| Ca (mg g<sup>-1</sup>) | 11 | 0.743 | 3.30 | 13.4 |
| Cu (mg g<sup>-1</sup>) | 10 | 0.354 | 0.00490 | 22.3 |
| Fe (mg g<sup>-1</sup>) | 3 | 0.135 | 0.0407 | 22.2 |
| K (mg g<sup>-1</sup>) | 12 | 0.548 | 3.13 | 20.2 |
| Mg (mg g<sup>-1</sup>) | 8 | 0.538 | 0.712 | 19.6 |
| Mn (mg g<sup>-1</sup>) | 9 | 0.327 | 0.171 | 20.8 |
| Na (mg g<sup>-1</sup>) | 11 | 0.461 | 0.519 | 22.1 |
| P (mg g<sup>-1</sup>) | 11 | 0.511 | 0.809 | 16.2 |
| Zn (mg g<sup>-1</sup>) | 10 | 0.428 | 0.0624 | 23.7 |

**Table S3:**
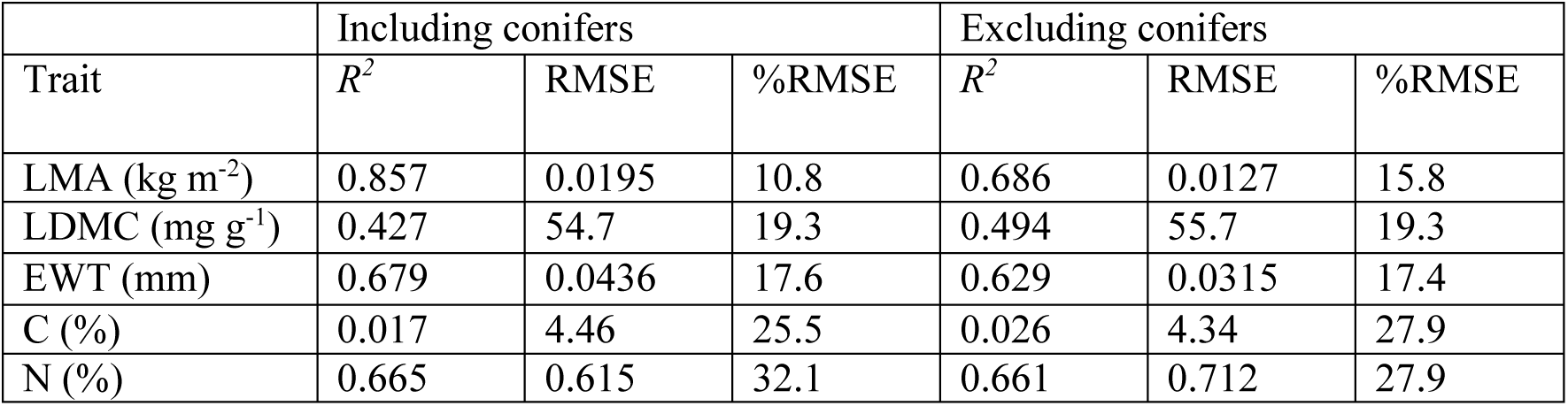
Summary statistics for external validation of pressed-leaf restricted-range PLSR models. The models were trained on CABO data (see Table S2) and applied to a dataset collected at Cedar Creek.

## Supplementary figures

**Fig. S1:**
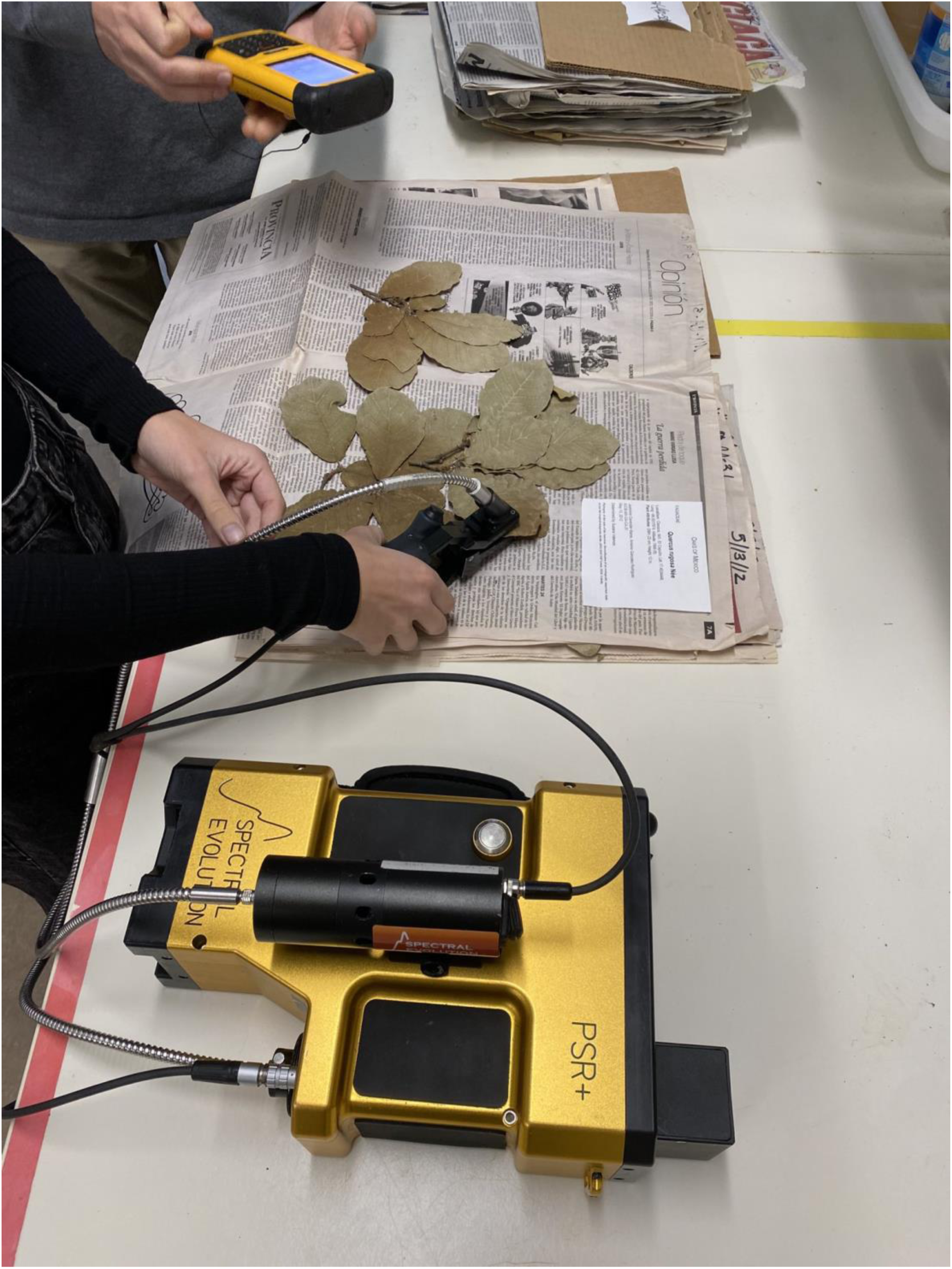
Two researchers (A. Scott and I. Clark) demonstrate a typical setup for measuring pressed-leaf spectra with a PSR+ 3500 spectroradiometer and leaf clip on unmounted pressed specimens. The specimen shown here was not part of this study.

**Fig. S2:**
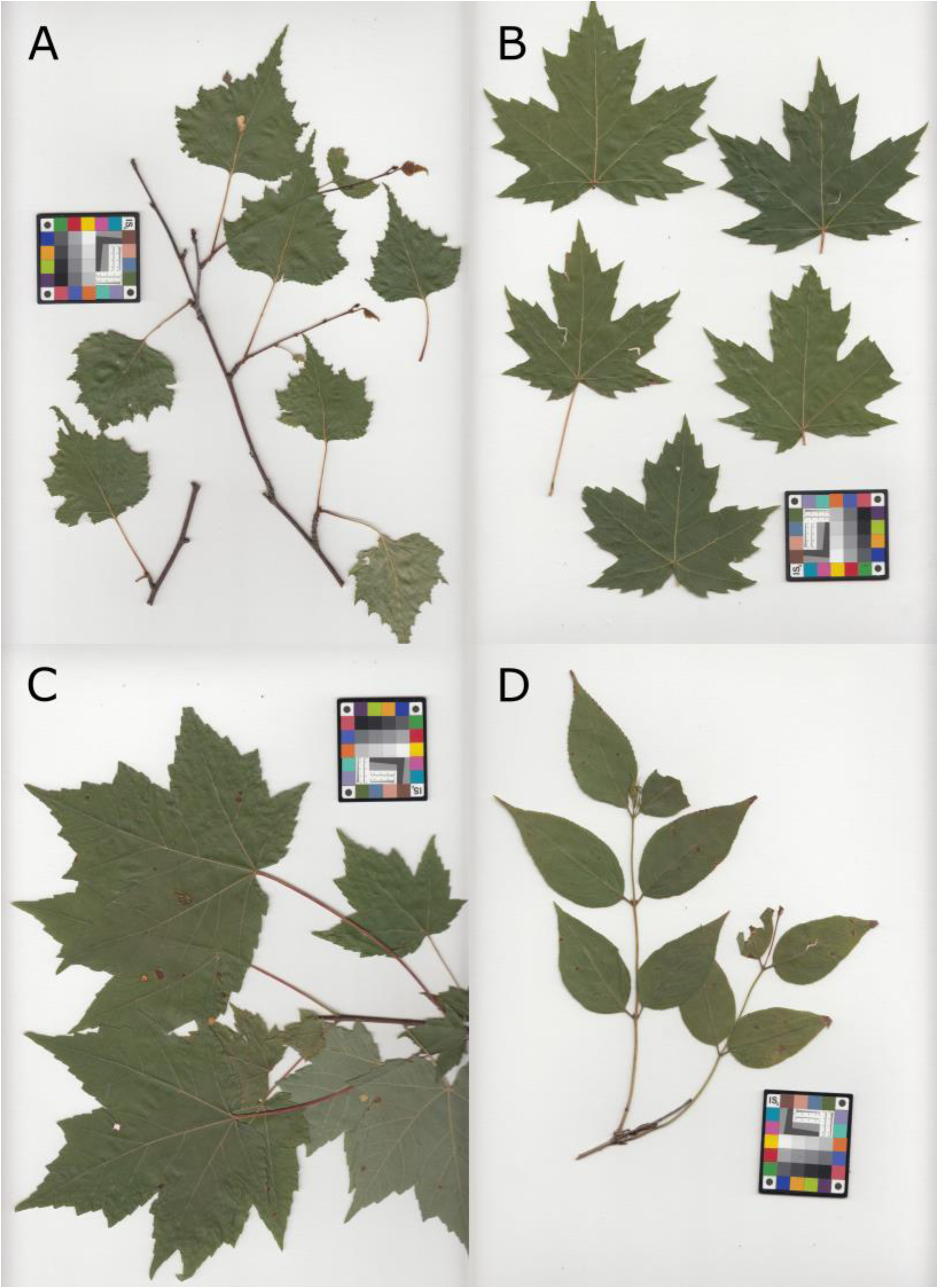
Scans of specimens that exemplify scores 0 and 1 on our discoloration scale. Specimens evaluated at 0 (A and B) showed no visible discoloration, as would be indicated by loss of green color (e.g. browning or blackening). Specimens evaluated at 1 (C and D) showed <10% discoloration throughout the leaf, or in some cases a more subtle, pervasive silvery or matte white appearance to the leaf surface in areas that remained otherwise green. On close inspection, these two samples both show some browning in a small portion of the total leaf area. Note that we do not consider spots or marks due to pathogen damage that were present at sampling time to be a form of discoloration; we also exclude some immature anthocyanic leaves as in (A). When in doubt, we examined scans of fresh leaves taken shortly after they were collected in the field. (Species: A, *Betula populifolia* Marshall; B, *Acer saccharinum* L.; C, *Acer rubrum* L.; D, *Diervilla lonicera* Miller.) All photos in Figs. S2-4 depict specimens included in the study, but were taken in March 2022, nearly three years after we took pressed-leaf spectral measurements. Due to continuing discoloration, their assigned scores as part of this set of example images may be larger than they were at the time we measured their spectra; the latter are what we used for data analysis.

**Fig. S3:**
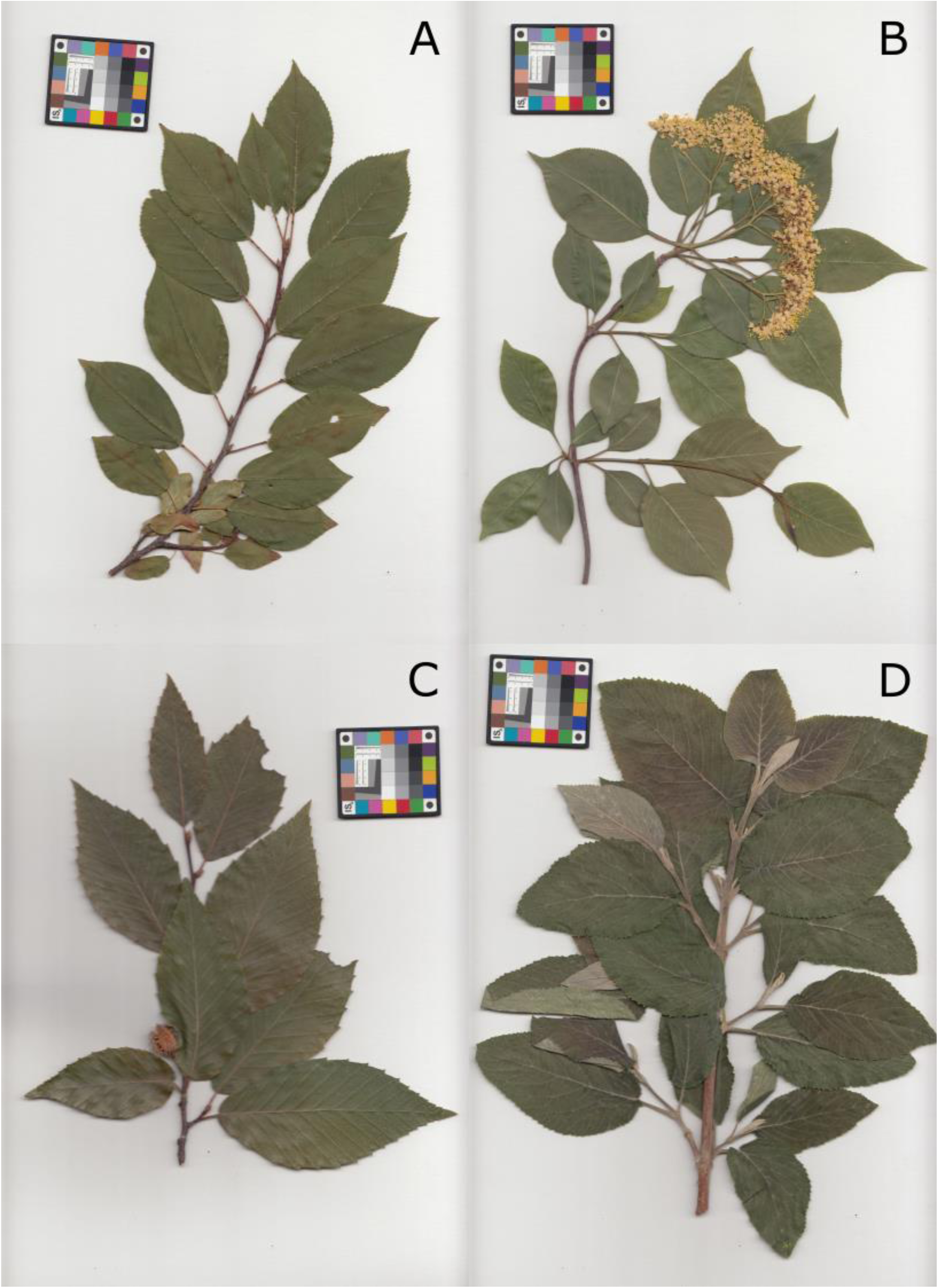
Scans of specimens that exemplify scores 2 and 3 on our discoloration scale. Specimens evaluated at 2 (A and B) showed 10-50% discoloration throughout the leaf. Here, A shows a number of dark streaks as well as more general browning on some leaves, while B shows more general darkening on some leaves. Specimens evaluated at 3 (C and D) showed 50-75% discoloration throughout the leaf. Here, C shows considerable browning across much of its leaf area, as well as a silvery appearance across much of the leaf surface. D shows considerable blackening across much of its leaf area. (Species: A, *Prunus virginiana* L.; B, *Viburnum lentago* L.; C, *Fagus grandifolia* Ehrhart; D, *Viburnum lantana* L.)

**Fig. S4:**
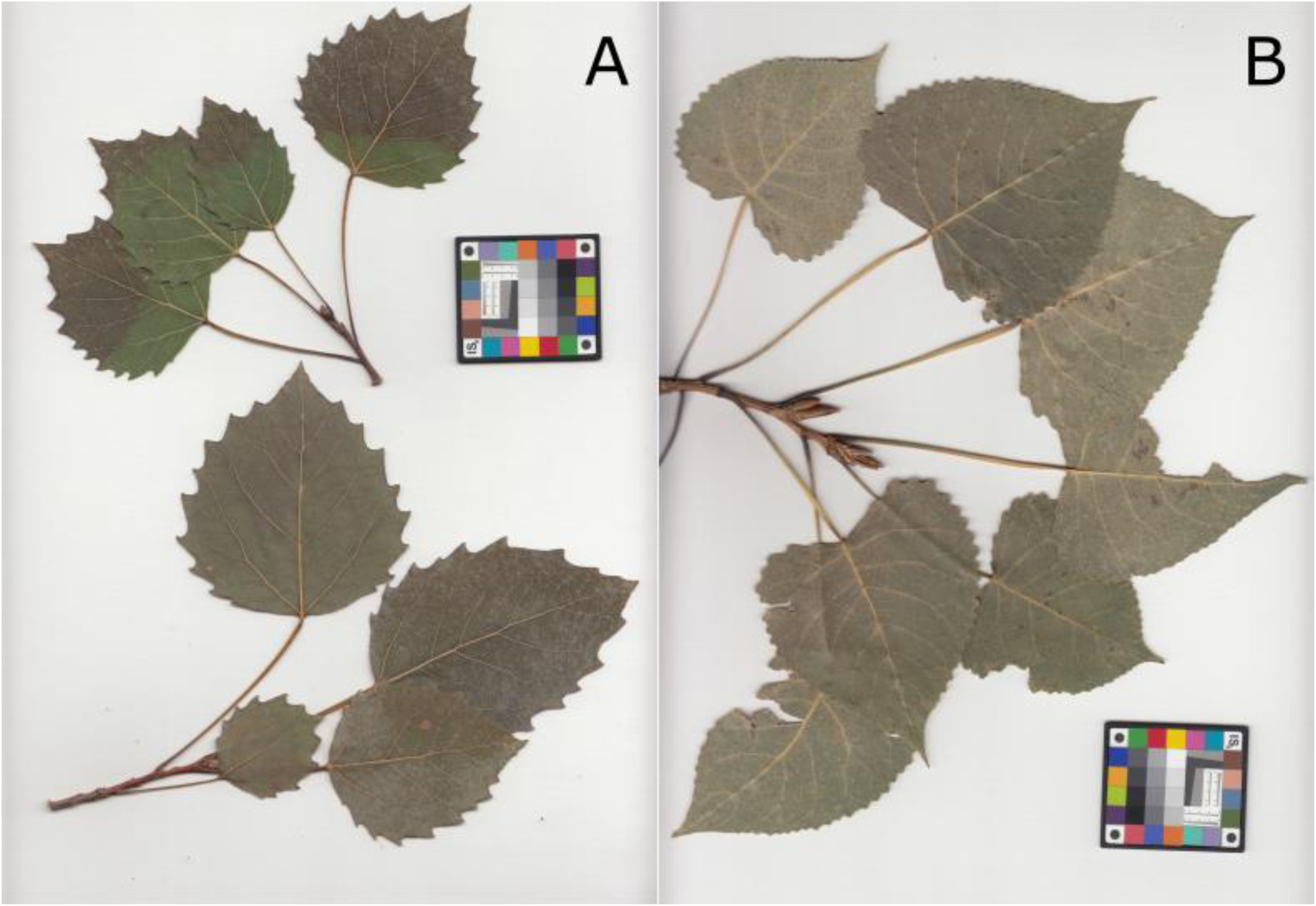
Scans of specimens that exemplify score 4 on our discoloration scale. These specimens showed nearly complete (>75%) discoloration, often making it difficult to take spectral measurements on a spot that retained green color. In some cases (A), the leaves retained patches of green color, while in others (B) the discoloration was nearly complete. (Species: A, *Populus grandidentata* Michaux; B, *Populus deltoides* W. Bartram ex Marshall.)

**Fig. S5:**
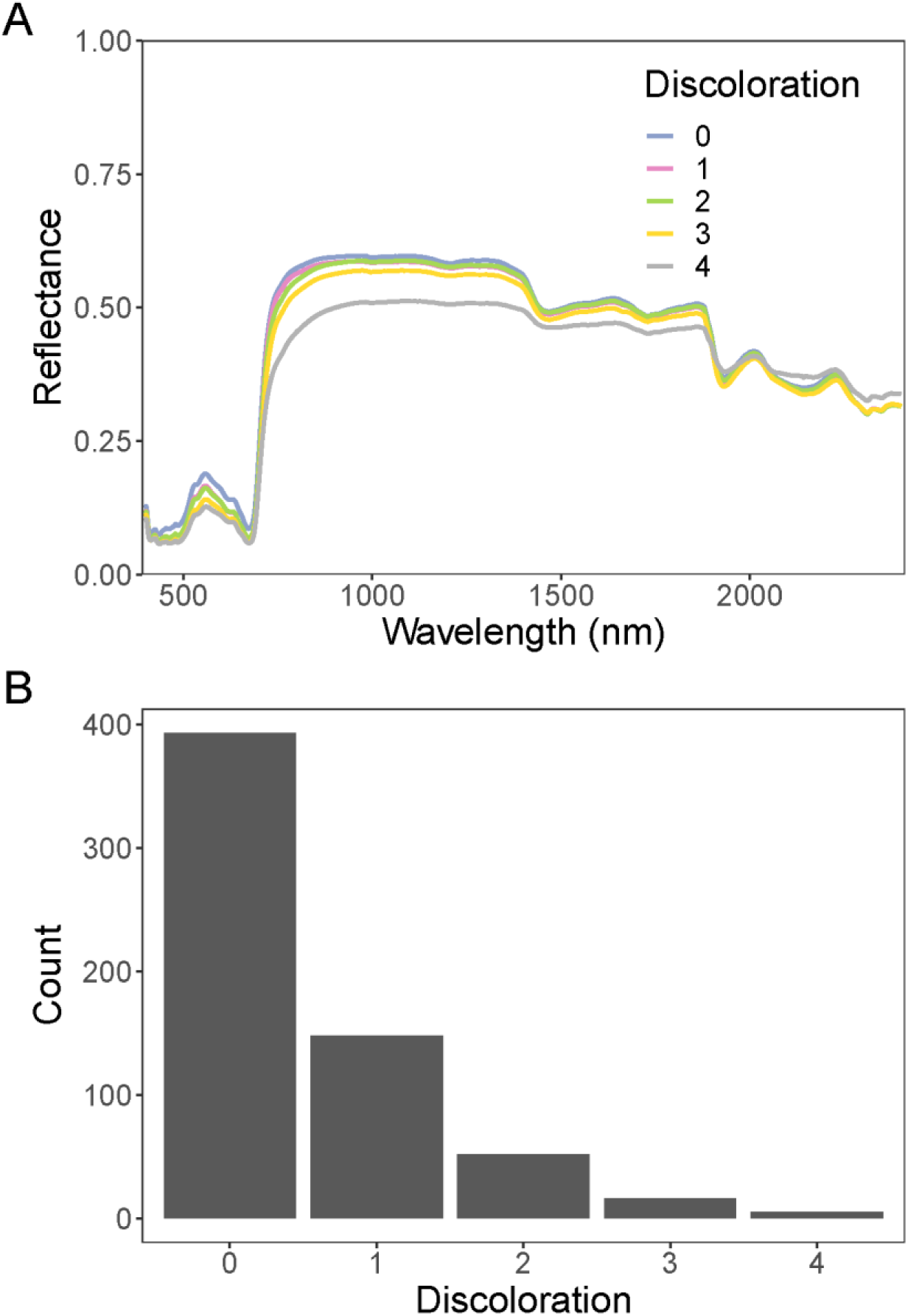
(A) The wavelength-wise mean reflectance spectrum of leaves at each discoloration score. In general, more discolored specimens have lower visible and NIR reflectance and a blunted red edge. This representation does not separate changes in spectra due to discoloration from aspects of spectral variation that might correlate with predisposition to discoloration. (B) A histogram of the number of pressed specimens evaluated at each discoloration stage at the time of spectral measurements.

**Fig. S6:**
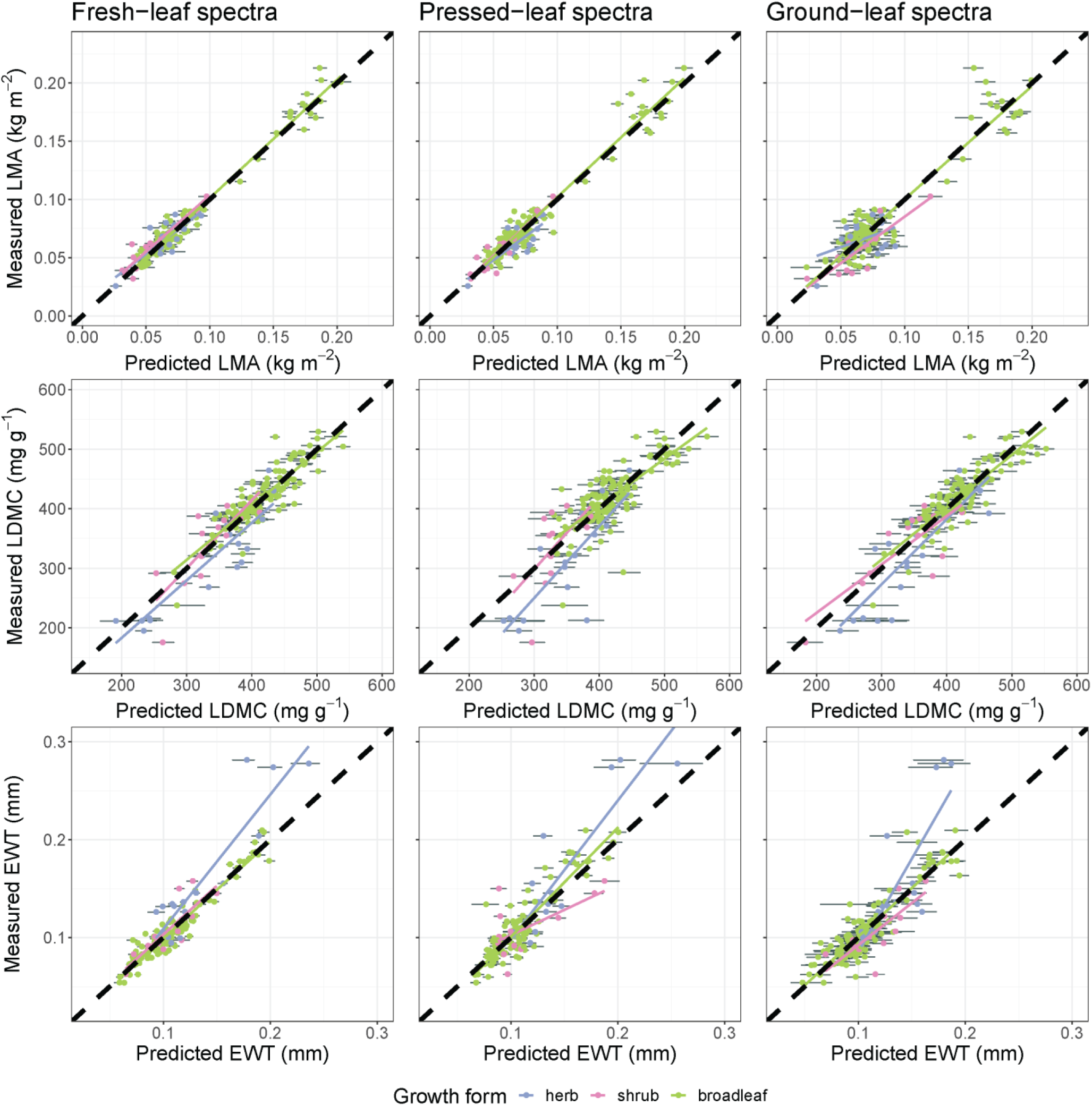
Internal validation results for predictions of LMA, LDMC, and EWT from fresh- (left), pressed- (middle), and ground-leaf (right) spectra. In each panel, each functional group has a separate ordinary least-squares regression line overlaid on top of the thick dashed 1:1 line. The error bars for each data point are 95% confidence intervals calculated from the distribution of predictions based on the ensemble of 100 models produced from jackknife analyses.

**Fig. S7:**
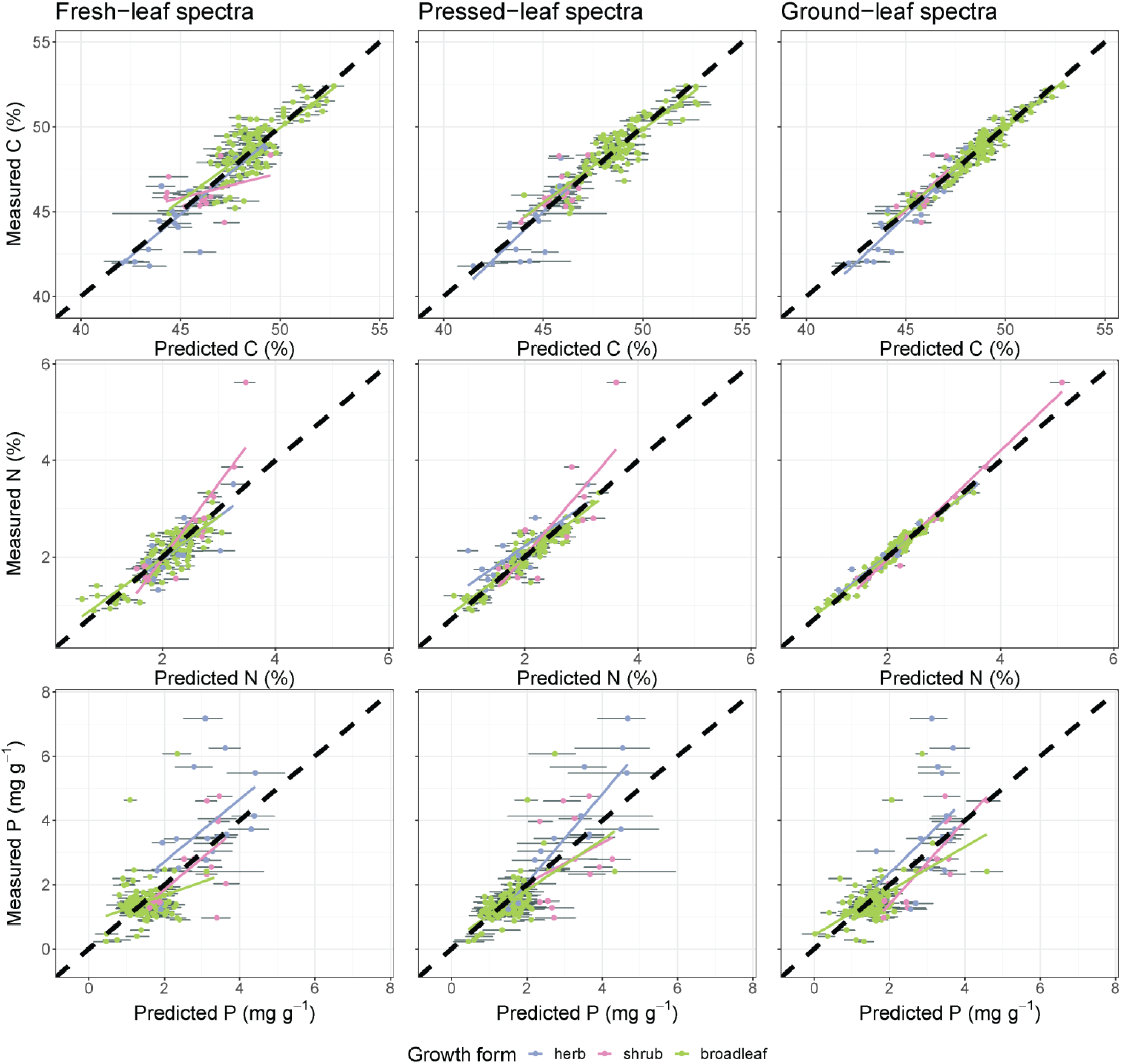
Internal validation results for predictions of C, N, and P from fresh- (left), pressed- (middle), and ground-leaf (right) spectra, displayed as in Fig. S6.

**Fig. S8:**
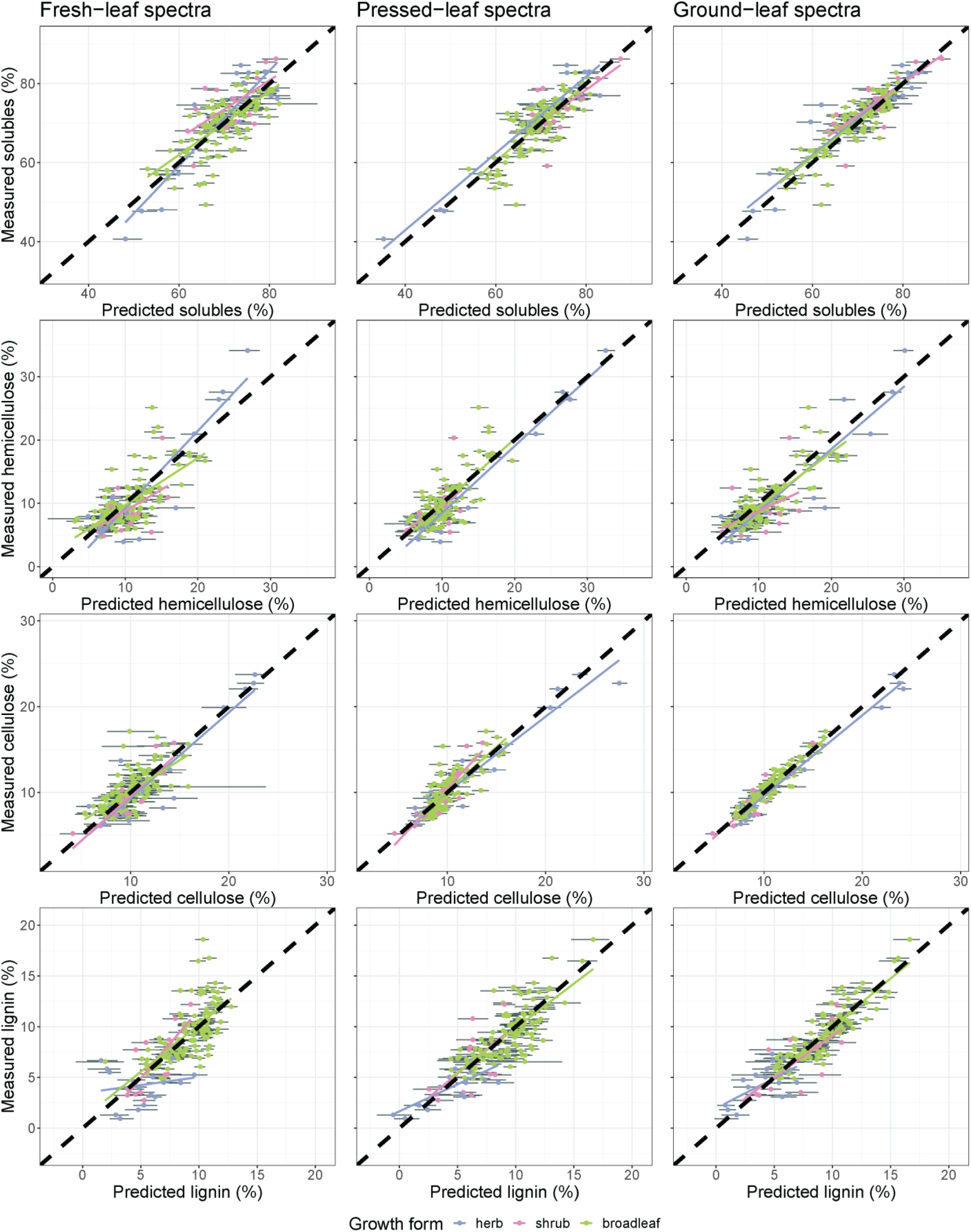
Internal validation results for predictions of soluble cell contents, hemicellulose, cellulose, and lignin from fresh- (left), pressed- (middle), and ground-leaf (right) spectra, displayed as in Fig. S6.

**Fig. S9:**
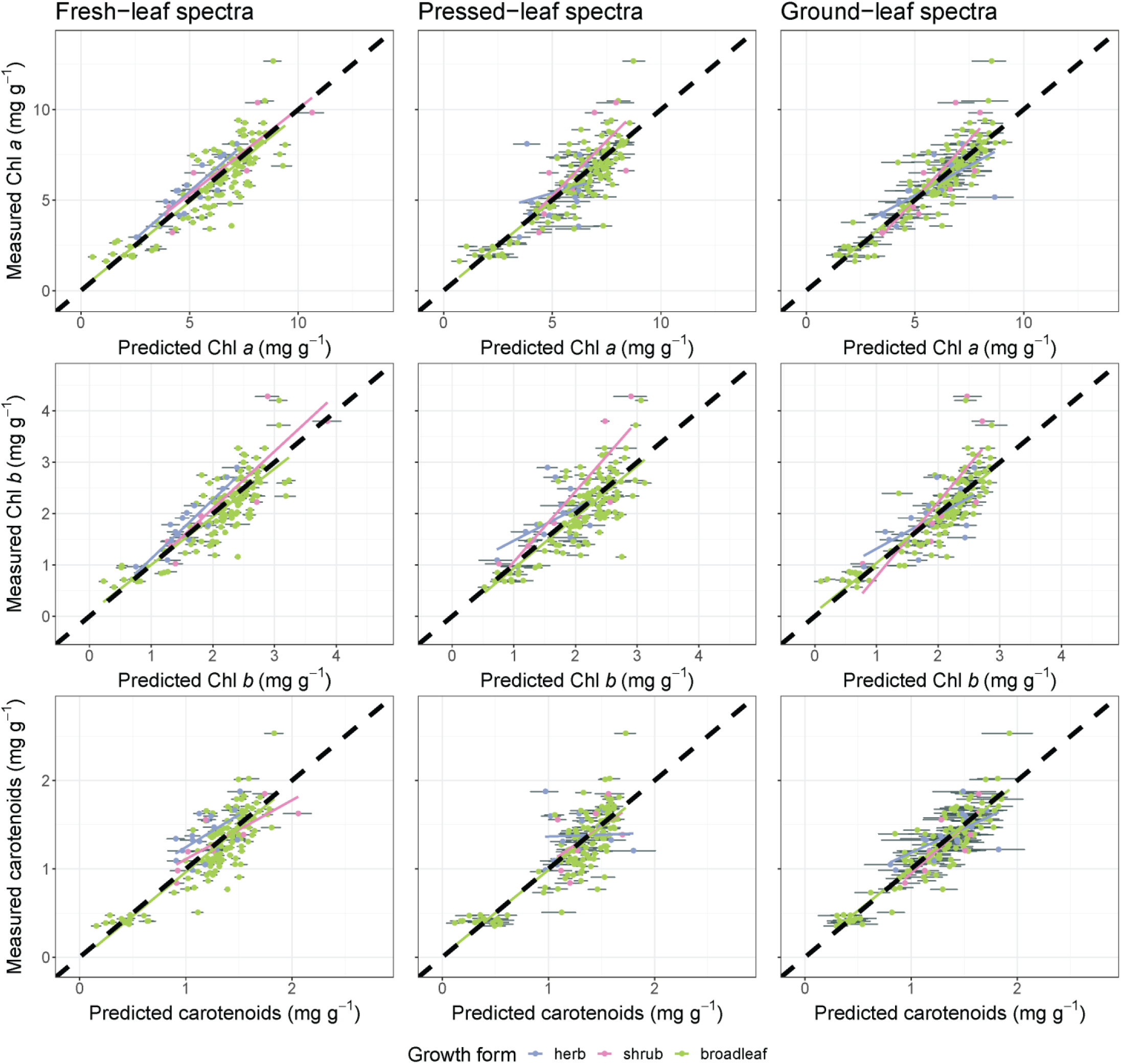
Internal validation results for predictions of chlorophyll *a*, chlorophyll *b*, and total carotenoids from fresh- (left), pressed- (middle), and ground-leaf (right) spectra, displayed as in Fig. S6.

**Fig. S10:**
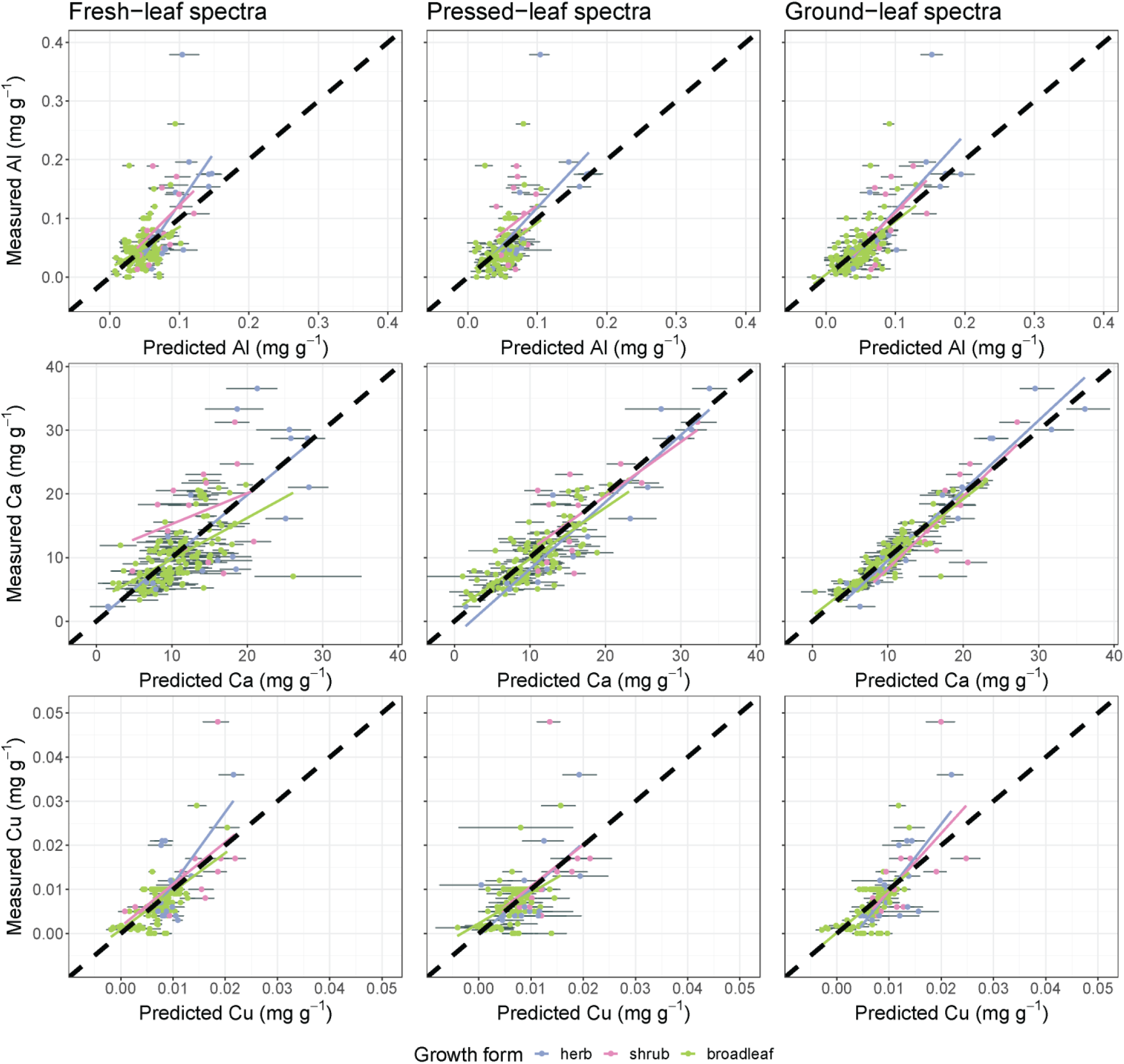
Internal validation results for predictions of Al, Ca, and Cu from fresh- (left), pressed- (middle), and ground-leaf (right) spectra, displayed as in Fig. S6.

**Fig. S11:**
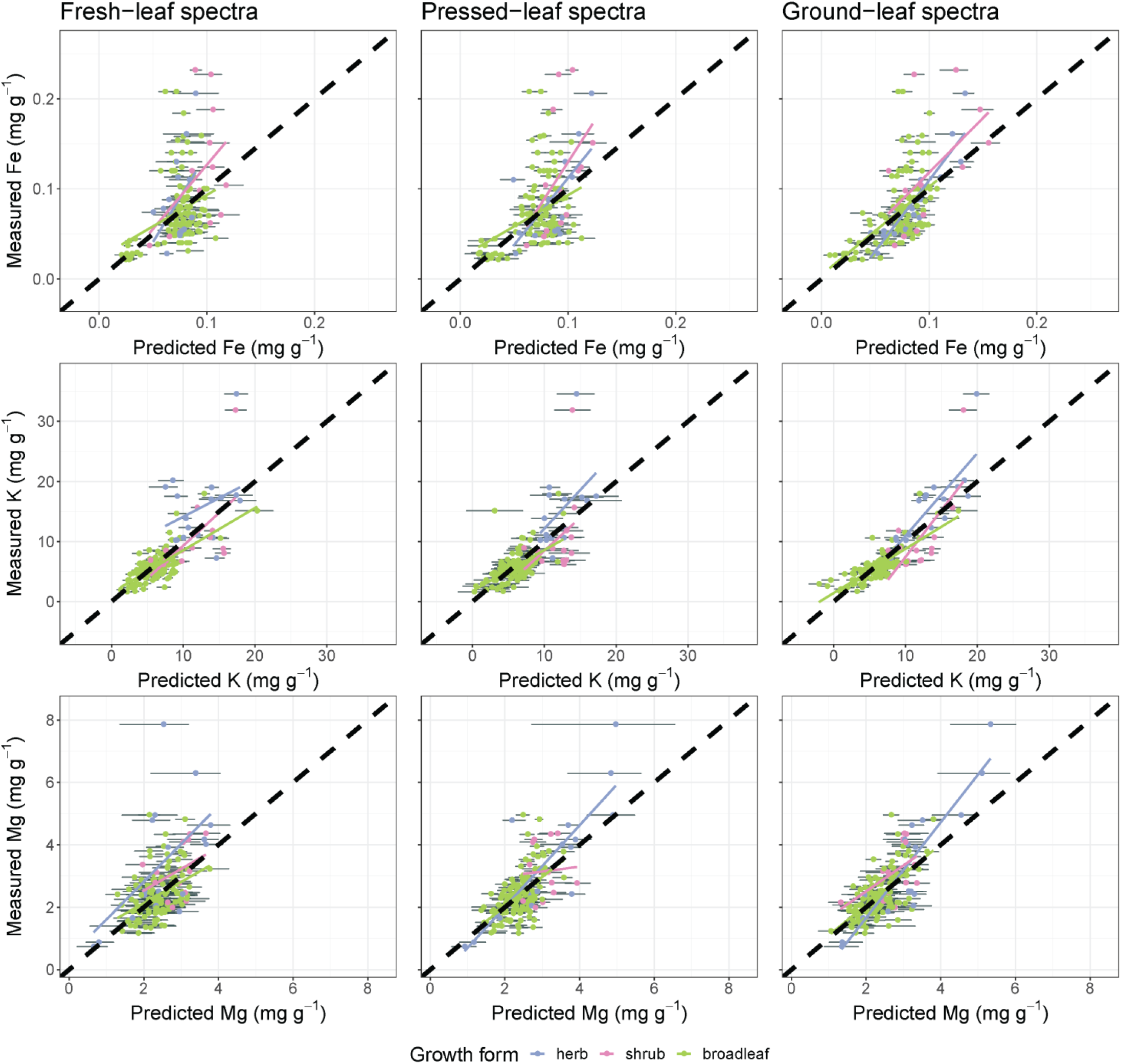
Internal validation results for predictions of Fe, K, and Mg from fresh- (left), pressed- (middle), and ground-leaf (right) spectra, displayed as in Fig. S6.

**Fig. S12:**
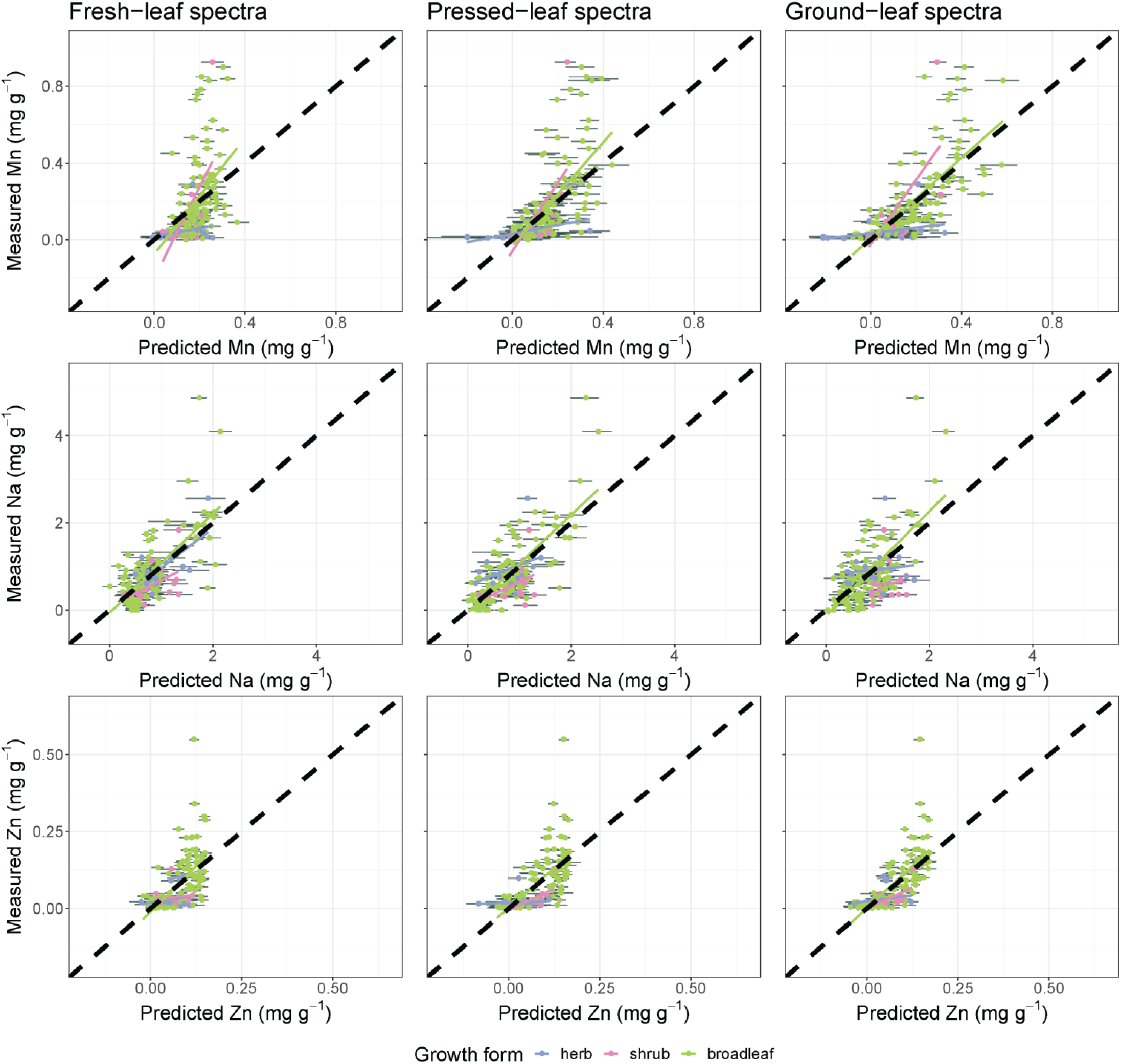
Internal validation results for predictions of Mn, Na, and Zn from fresh- (left), pressed- (middle), and ground- leaf (right) spectra, displayed as in Fig. S6.

**Fig. S13:**
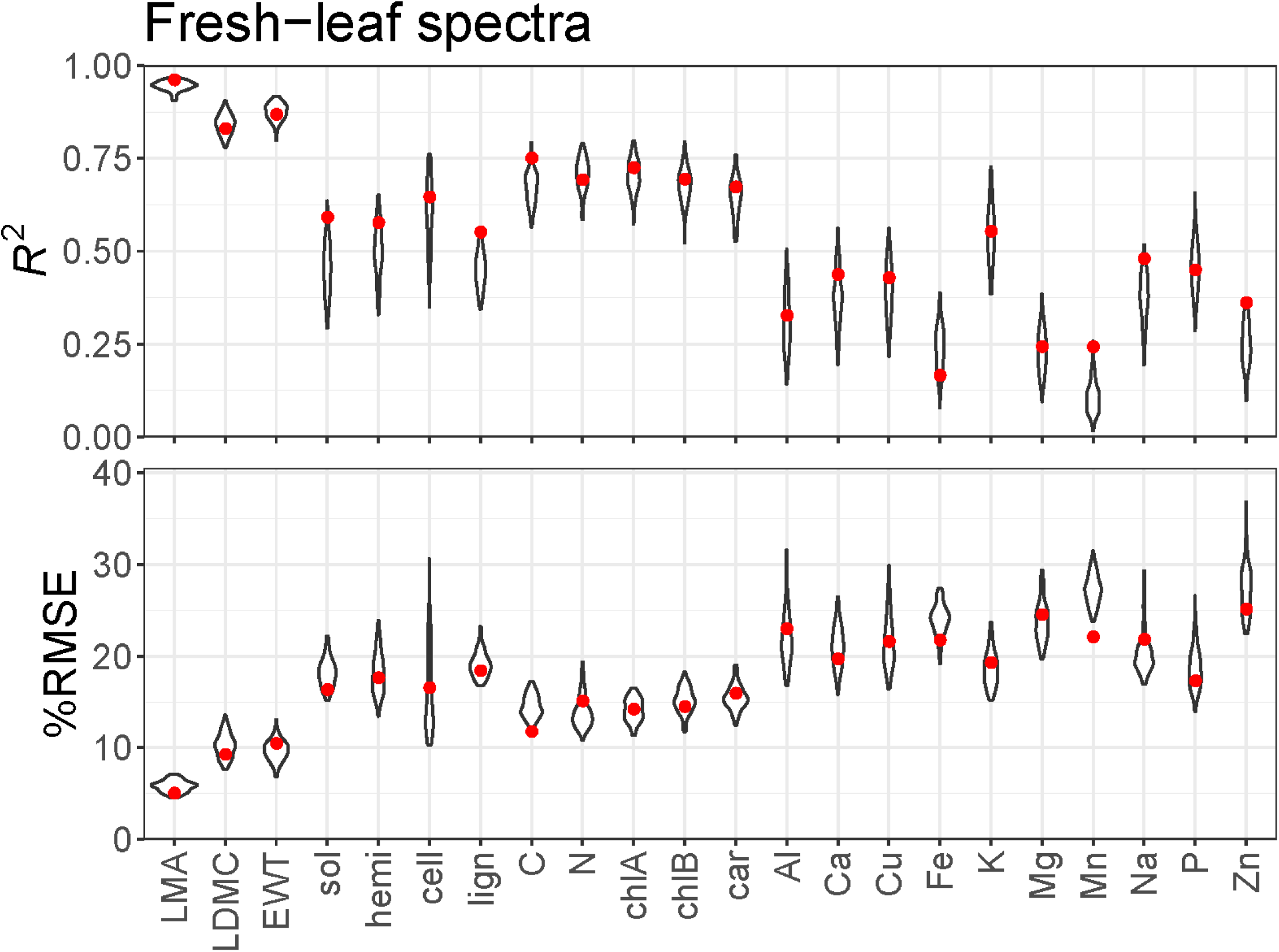
The distributions of fresh-leaf spectral model performance statistics for each trait based on 100 jackknife resamples from the calibration data set. The red dots show the *R^2^* and %RMSE from applying the ensemble of models to the internal validation subset. Abbreviations: sol = solubles, hemi = hemicellulose, cell = cellulose, lign = lignin, car = total carotenoids.

**Fig. S14:**
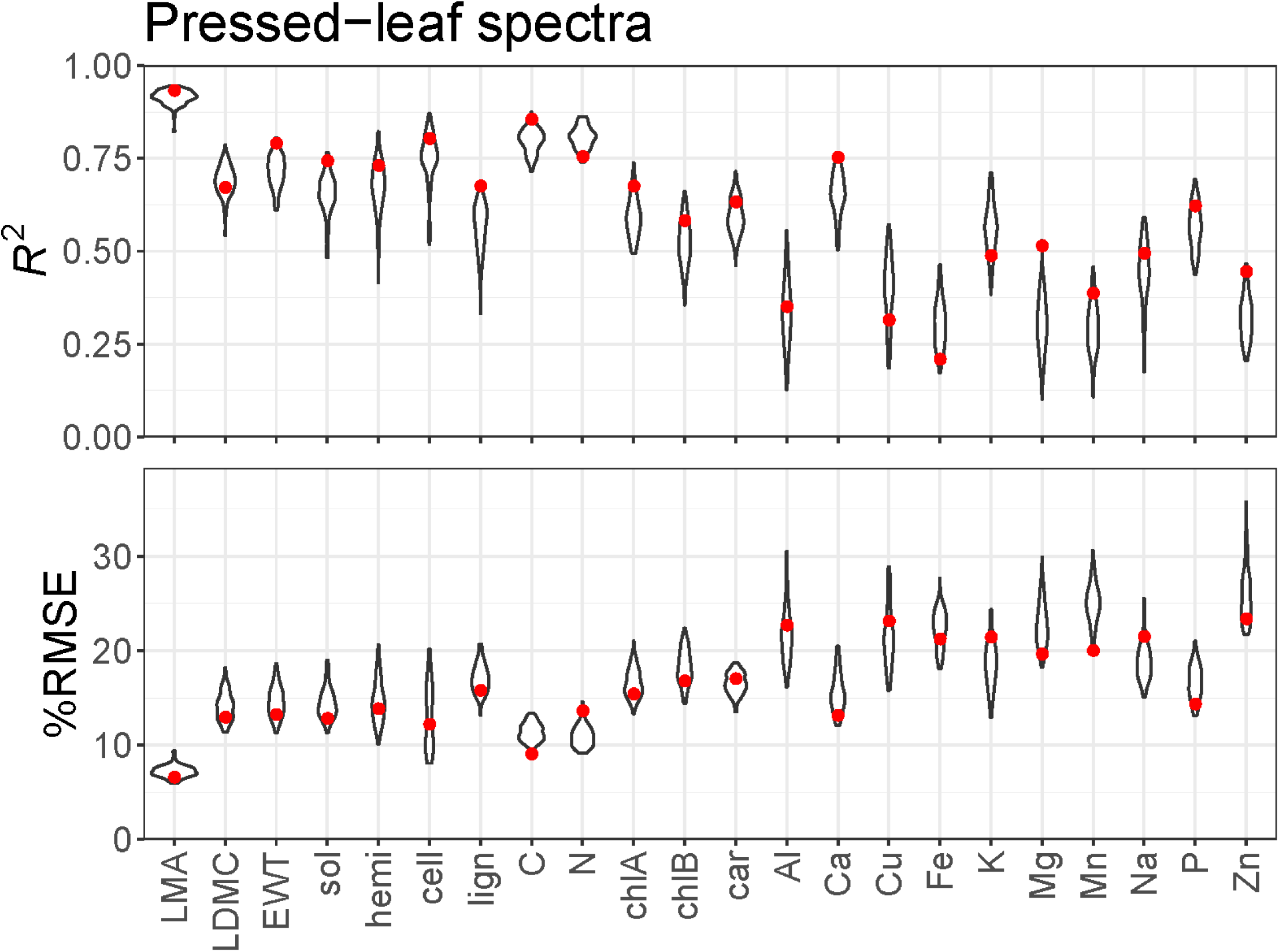
The distributions of pressed-leaf spectral model performance statistics for each trait based on 100 jackknife resamples from the calibration data set. The red dots show the *R^2^* and %RMSE from applying the ensemble of models to the internal validation subset. Abbreviations: sol = solubles, hemi = hemicellulose, cell = cellulose, lign = lignin, car = total carotenoids.

**Fig. S15:**
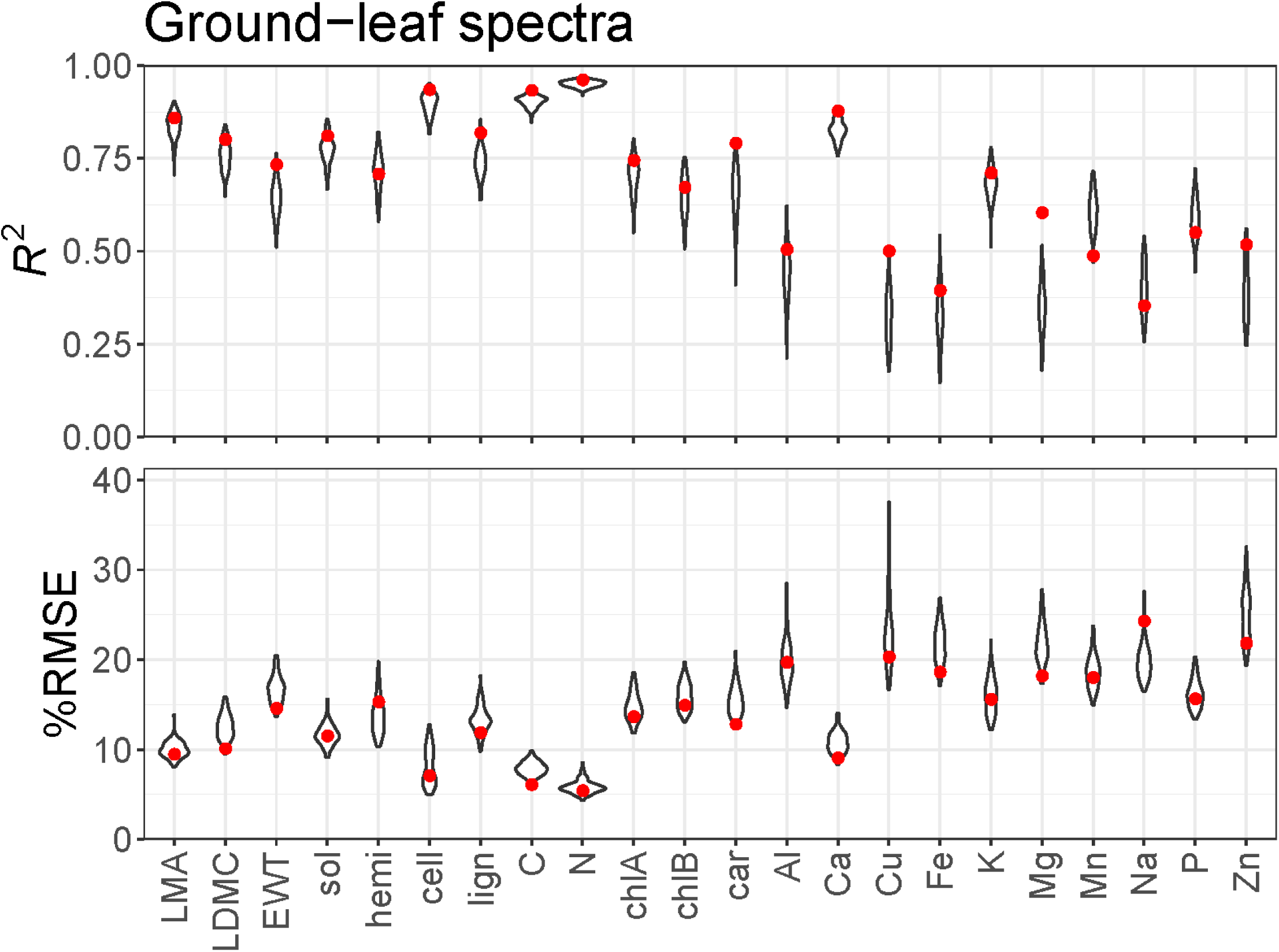
The distributions of ground-leaf spectral model performance statistics for each trait based on 100 jackknife resamples from the calibration data set. The red dots show the *R^2^* and %RMSE from applying the ensemble of models to the internal validation subset. Abbreviations: sol = solubles, hemi = hemicellulose, cell = cellulose, lign = lignin, car = total carotenoids.

**Fig. S16:**
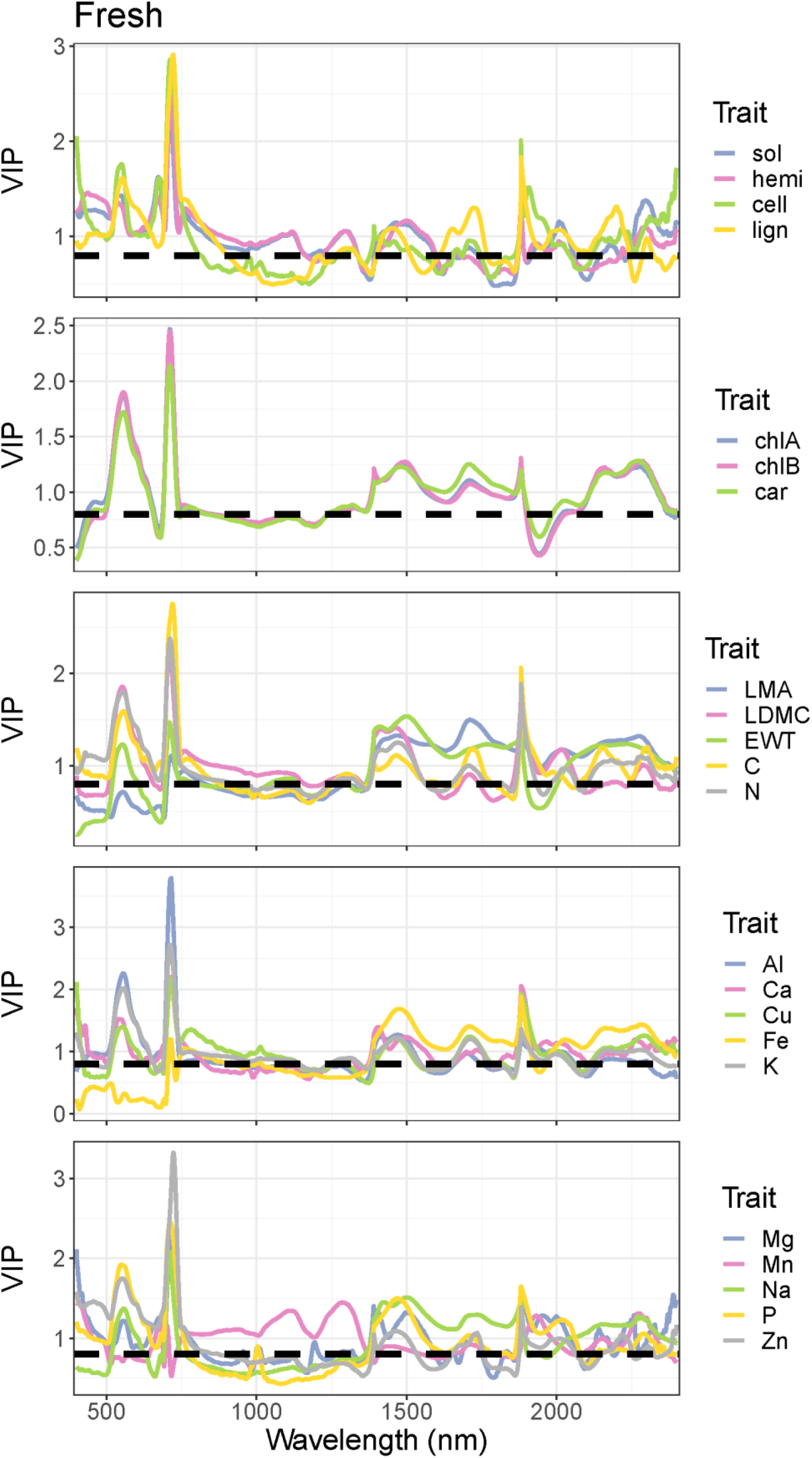
The variable importance of projection (VIP) metric calculated for all traits from fresh-leaf spectra. Abbreviations: sol = solubles, hemi = hemicellulose, cell = cellulose, lign = lignin, car = total carotenoids. The dashed horizontal line at 0.8 represents a heuristic threshold for importance suggested by Burnett et al. (2021).

**Fig. S17:**
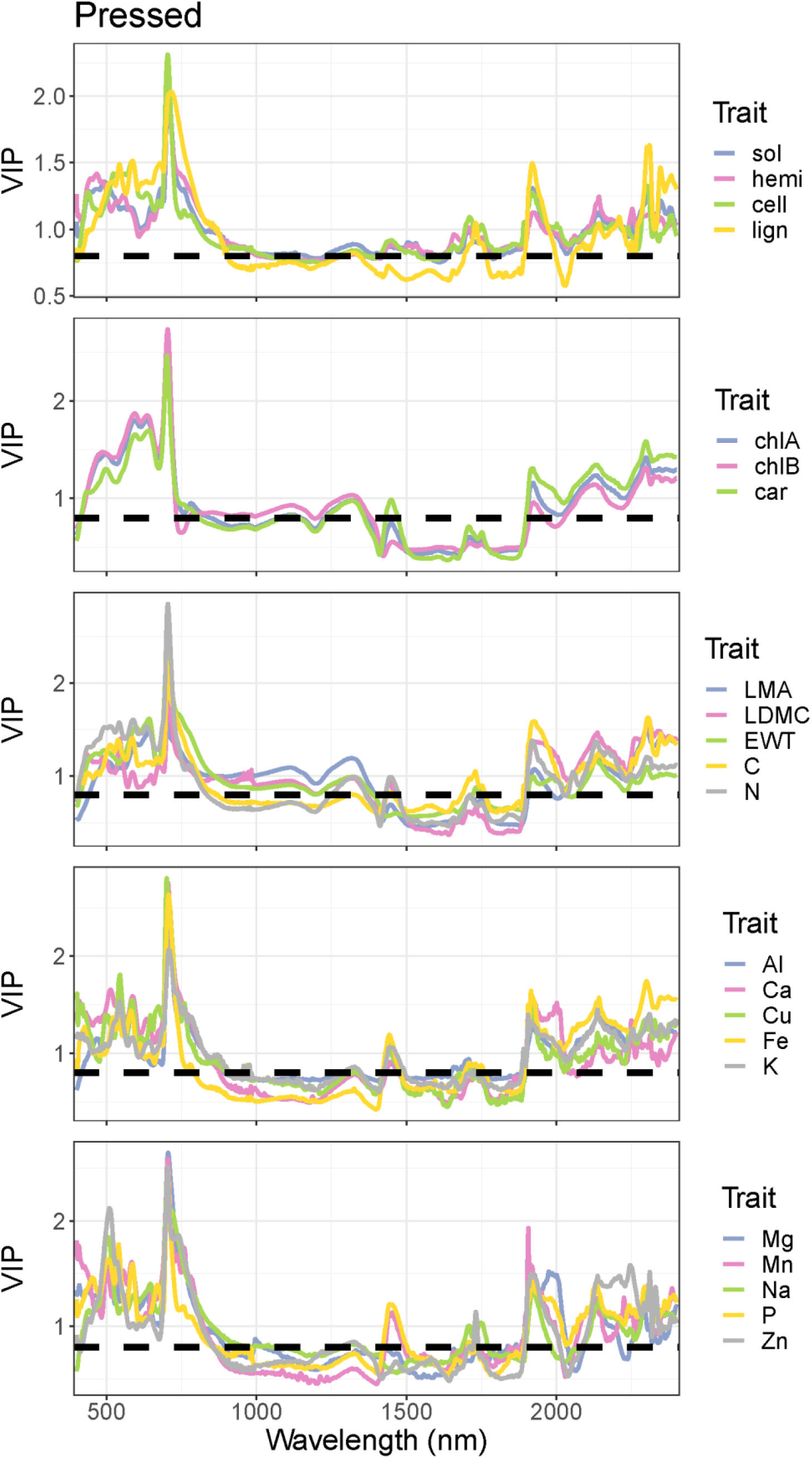
The variable importance of projection (VIP) metric calculated for all traits from pressed-leaf spectra. Abbreviations: sol = solubles, hemi = hemicellulose, cell = cellulose, lign = lignin, car = total carotenoids. The dashed horizontal line at 0.8 represents a heuristic threshold for importance suggested by Burnett et al. (2021).

**Fig. S18:**
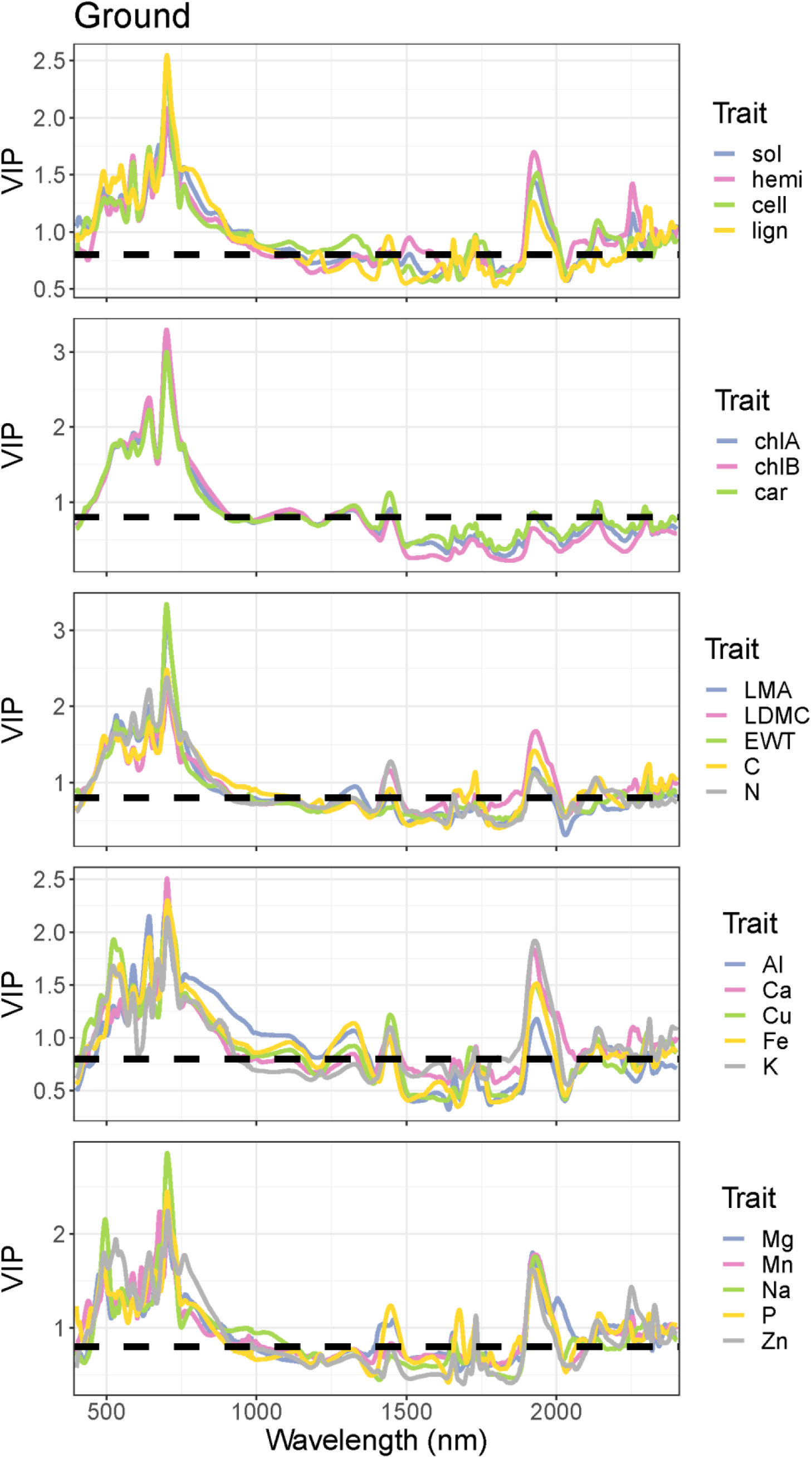
The variable importance of projection (VIP) metric calculated for all traits from ground-leaf spectra. Abbreviations: sol = solubles, hemi = hemicellulose, cell = cellulose, lign = lignin, car = total carotenoids. The dashed horizontal line at 0.8 represents a heuristic threshold for importance suggested by Burnett et al. (2021).

**Fig. S19:**
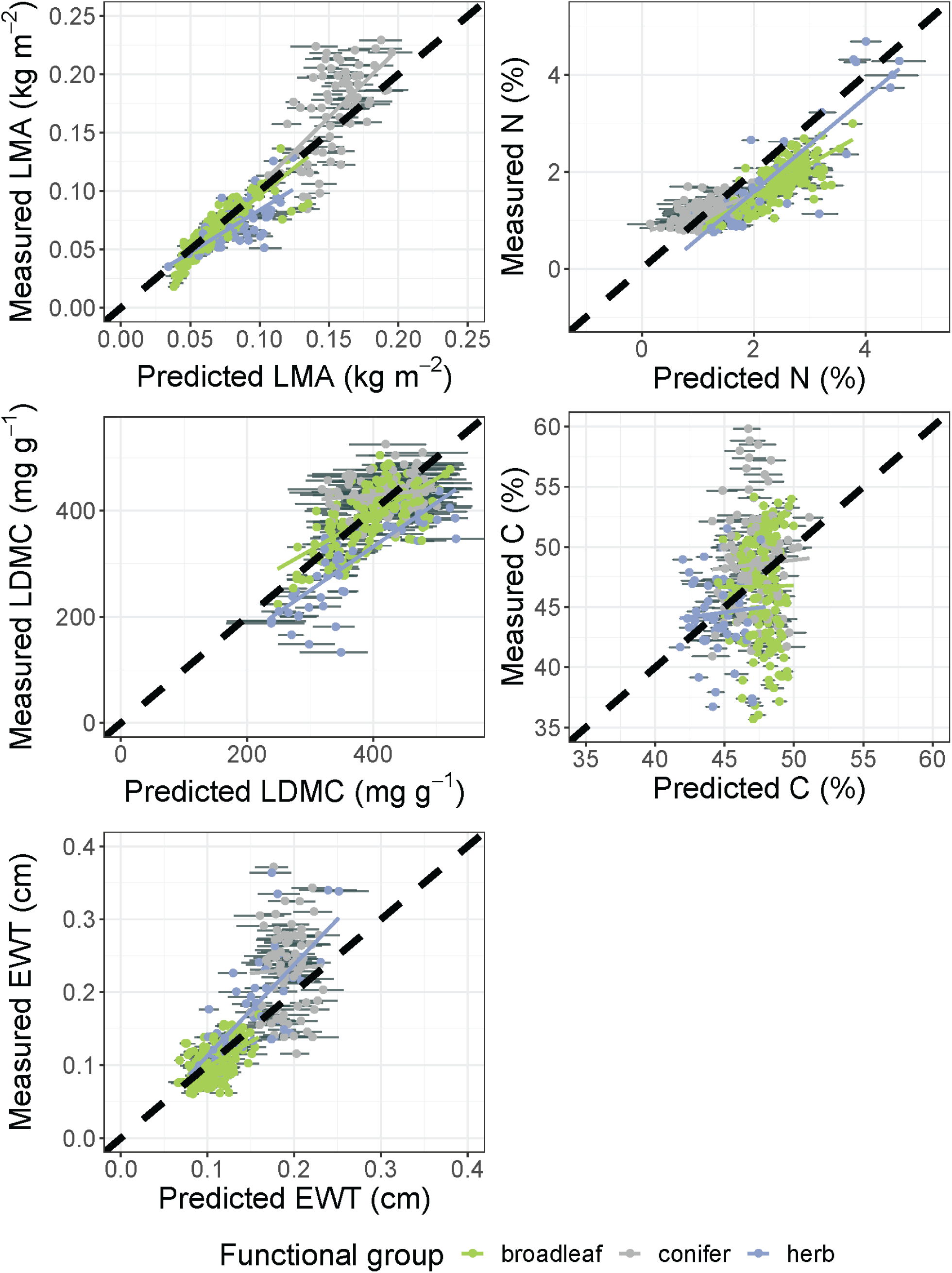
External validation results for predictions of various traits in Cedar Creek data based on models trained on CABO data, restricted to 1300-2500 nm. Each panel is displayed as in Figs. S6-12.

## Notes

### Competing Interest Statement

The authors have declared no competing interest.

### Summary of Updates

Many changes; new figures/table, text reorganized

